# Predicting specificity of TCR-pMHC interactions using machine learning and biophysical models

**DOI:** 10.1101/2025.04.04.647165

**Authors:** Martin Culka, Nicolas W. Lounsbury, William Thrift, Santrupti Nerli, Andrew Wallace, Gergő Nikolényi, Darya Orlova, Kiran Mukhyala, Mohammed AlQuraishi

## Abstract

Understanding the mechanism of T-cell activation and T-cell receptor (TCR) discrimination of MHC-presented epitope peptides (pMHCs) remains an open problem. Machine learning (ML)-based prediction of TCR specificity has gained considerable recent attention. However, the capacity of current models to generalize to peptides unseen during training is currently unknown. Here, we use a proprietary cancer-patient data set that profiles TCR binding to novel regions of peptide space to show that peptide generalization remains an unsolved problem. Specifically, we show that while ML methods have demonstrable utility in predicting TCR specificity for known peptides, they fail to generalize to novel peptides. We also show that physics-based methods utilizing classical energy functions outperform ML methods when predicting TCR binding to novel peptides but underperform them on known peptides. In light of these observations, we develop a new ML model that leverages general knowledge acquired by protein foundation models to achieve better or comparable performance than either ML or biophysical methods on both in- and out-of-distribution TCR-pMHC specificity prediction. We furthermore analyze model performance as a function of distance of TCR sequence specificity between the training and test sets to quantitatively characterize the generalization potential of any given TCR-pMHC model. Our analysis sheds light on the status of modeling TCR-pMHC interactions and suggests new paths forward for continued method development and data acquisition.

---

T cells play a crucial role in immunosurveillance^1^, yet how their receptors (TCRs) respond to antigenic peptides presented by the major histocompatibility complex (pMHC) is not fully understood. A better understanding of how T cells operate would inform the development of therapeutic strategies for autoimmune diseases, cancer immunotherapy, and vaccine design, where modulation of T cell responses is critical. Without mechanistic understanding, efficient and reliable selection of a pMHC-specific TCR from a TCR repertoire (>10^8^ TCRs)^2^ remains highly challenging.

The existence of high-throughput TCR-pMHC binding data has generated interest in building machine learning (ML) models of TCR specificity. Numerous approaches have been proposed to date, including ones based on convolutional neural networks^3,4^, meta-learning^5^, or transformers^6–8^ (see recent reviews^9–12^ and Table S1 for a more comprehensive overview). These models are often trained directly on TCR-pMHC pairings using raw amino acid sequences as the input and thus do not leverage general purpose knowledge acquired by protein foundation models^13,14^, although a subset does first perform unsupervised pre-training on unlabeled data^7^. Given the limited nature of publicly available data, most ML models use only complementarity-determining regions (CDRs), specifically CDR3*β* sequences, to represent TCRs and peptide sequences to represent pMHCs.

To achieve full utility, ML models must generalize to previously unseen peptides. The degree to which current data sets sufficiently cover pMHC and TCR space to permit such generalization remains unclear, however. So far, ML models have demonstrated promising results in applications where peptide sequences do not vary considerably between training and test sets, even when only limited amounts of training data are available^4^. Testing generalization to new peptide sequences is challenging however, given the limited amounts of publicly available TCR-pMHC pairing data. Furthermore, some model assessments suffer from poor experimental design. For instance, multiple different strategies exist for generating non-productive (negative) pairings. It has been shown that using TCRs from healthy patients as negatives leads to much more optimistic results relative to shuffling epitopes and TCRs within the same data set, despite the fact that the latter design more closely resembles real-world deployment scenarios^8,15^. Ideally, standardized blind testing procedures would be performed for TCR-pMHC modeling, similar to the CASP assessments for protein structure prediction^16^. Some nascent efforts have taken place, including the peptide-specific IMMREP_2022^17^ and panpeptide IMMREP23^18^ challenges. The former, however, does not test generalization to novel peptides, while the latter exhibits overlap between training and test sets with respect to both epitopes and TCRs, compromising the integrity of the experiment. Additionally, both challenges used publicly available data biased toward a limited set of predominantly viral peptides. The field thus remains lacking in rigorous testing standards, especially with respect to practical use cases such as in cancer treatment.

Here, we aim to rectify these shortcomings through a series of analyses of TCR-pMHC specificity prediction methods, including supervised ML models and non-supervised structure- and physics-based scoring functions (such as protein-protein docking scores or force field interaction energies). We also develop and assess a class of new ML methods, TCRcube, which leverage general protein knowledge acquired by protein foundation models (ESM2^13^ and AlphaFold2^19^) and model full-length TCR sequences. We first focus on the peptide-specific task (unseen TCR or in-distribution generalization) and show that TCRcube models are either on par with or outperform state-of-the-art ML approaches on the standardized IMMREP_2022 benchmark set. All ML methods in turn outperform non-supervised scoring functions on this task. Next, we tackle out-of-distribution generalization by testing pan-peptide models on unseen epitope peptides and TCRs. We show that TCRcube models achieve comparable performance to non-supervised scoring functions, which outperform existing ML methods on out-of-distribution tasks. In particular, we find that ML methods claiming performant out-of-distribution generalization fail to generalize, with previously promising results likely due to problematic experimental design. To unambiguously test this hypothesis, we conduct a zero-shot test using proprietary data collected from cancer patients by Genentech and Adaptive Biotechnologies. Since the majority of publicly available data is on viral epitopes, the problem of predicting cancer epitopes (neoantigens) is a highly challenging generalization task for TCR-pMHC models. We show that all models, including TCRcube, struggle to make useful predictions in this zero-shot setting. We employ TCRcube to better understand why peptide-specific generalization is tractable by current ML approaches while out-of-distribution generalization is not, and provide an actionable blueprint for the design of systematic data acquisition experiments that can bridge the generalization gap.

## Results

### TCRcube model architecture and training procedure

The vast majority of publicly available TCR-pMHC binding data is binary, indicating whether a specific pMHC is paired with a specific TCR without quantification of binding affinity. This reality informs the design of the TCRcube family of models, which we formulate as binary classifiers. Approximately 13,000 binary binding measurements exist, which we use as the basis for model training; further details about data set construction and splitting strategies are described in methods and subsequent sections. TCRcube models (Figure 1A) accept as input paired TCR-pMHC sequences represented either by raw amino acid identities or latent embeddings from protein foundation models. The latter category includes the sequenced-based ESM2 protein language model and AlphaFold2-based (AF2) representations extracted from its evoformer or structure modules. TCRcube models begin with fully-connected neural networks that project the input representations of peptide, MHC, and TCR into a common latent space. An outer product is then computed between these latent representations to form a 3rd-order tensor, whose entries are residue triplets. From each triplet, an energy term is computed using a ReLU activation function, and the resulting energies are summed and transformed into a binding probability of the TCR-pMHC pair. We use a three-body formulation to more faithfully describe the mechanics of peptide-MHC complex recognition by TCRs, by accounting for peptide-TCR interactions, as the majority of existing models do, as well as peptide-assisted TCR-MHC interactions and MHC-assisted peptide-TCR interactions.

**Figure 1.**
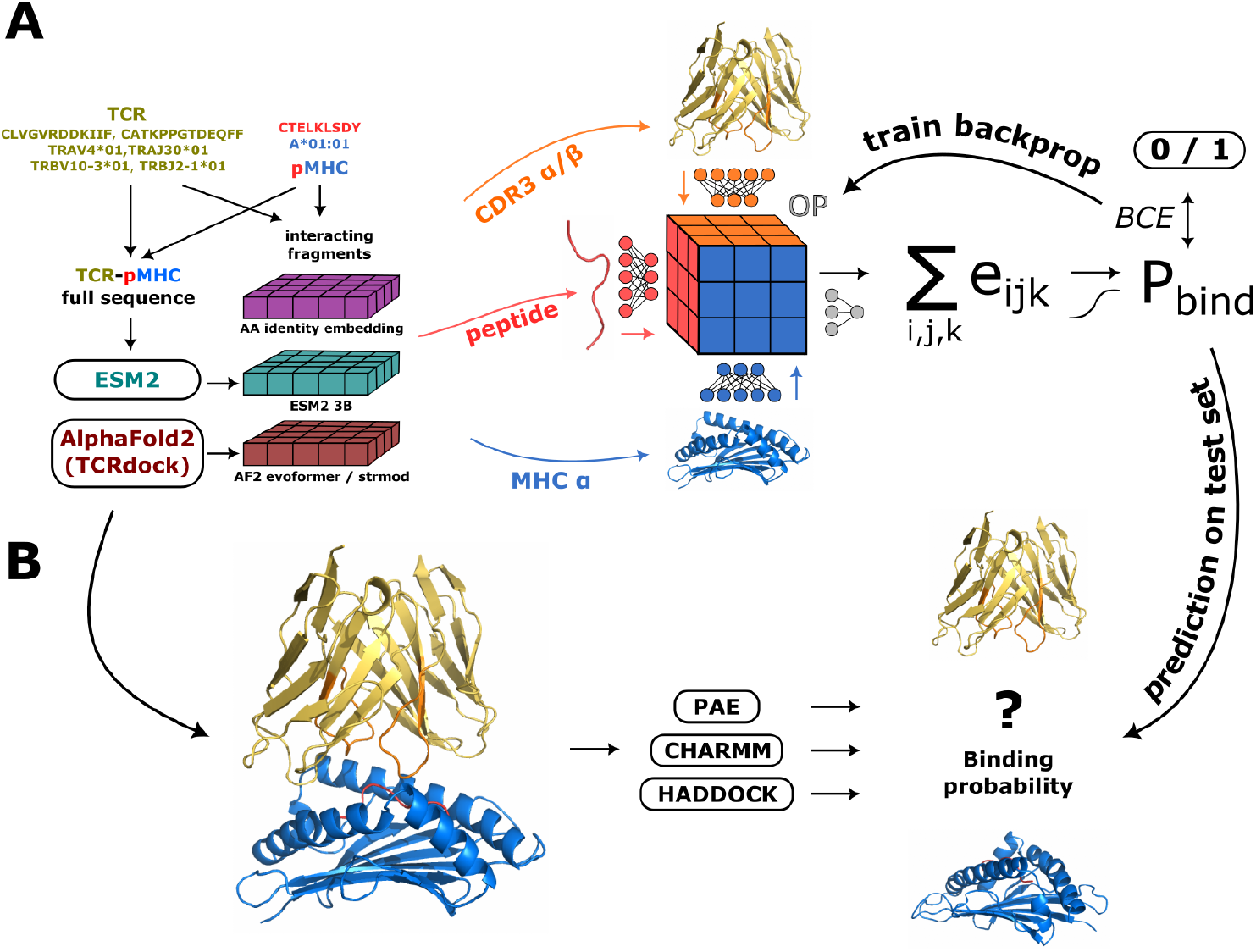
(**A)** Architecture of the TCRcube family of models. Putative TCR-pMHC complexes are first disassembled into constituent MHC, peptide, and TCR CDR3 loops using one of many possible representations (raw amino acid identity, ESM2, or AF2 embeddings). Linear projections of the three TCR-pMHC components (CDR3 sequence of TCR, peptide, and MHC) are then combined through an outer product (OP) to form a 3rd-order tensor. Tensor entries are transformed through an element-wise linear layer into energy terms e_ijk_. Total energy is finally computed by summing all energy terms and converted into a binding probability P_bind_. (**B)** Structure-based scoring functions are used as an alternative mode of binding probability prediction.

### Structure-based scoring function prediction

To complement ML-based methods, we employ structurally- and biophysically-grounded scoring functions to predict affinities of TCR-pMHC complexes (Figure 1B). For input, we use *in silico* structures of TCR-pMHC complexes predicted by the AF2-based TCRdock^20^ pipeline. Predicted complexes (of both positive and negative pairings) generally show reasonable binding geometry but with occasional clashes at the interface (in ∼30% of complexes, the distance between some heavy atoms at the TCR-pMHC interface is <1.5Å; see Figure S1). The simplest scoring function we consider is the PAE value outputted by AF2, which summarizes AF2’s own confidence in the prediction. We also employ the CHARMM^21^ force field, which we use to calculate interaction energy as well as correct clashes via energy minimization. The third and final approach we use redocks the predicted TCR-pMHC complex by first separating the pMHC from the TCR chain then uses HADDOCK^22^ to reform the complex. HADDOCK computes a new complex geometry and assigns it a score. We constrain HADDOCK docking by specifying which residues lie at the interface based on the original AF2-predicted TCR-pMHC complex.

### Peptide-specific evaluation of in-distribution generalization

We begin our assessment with a comparatively easy task, that of predicting TCR-pMHC binding of epitopes present in the training set. This is a problem on which existing ML models have demonstrated high accuracy. We use the IMMREP_2022 data set^17^, which organizes binding data into 17 categories based on epitope sequence (14,719 training and 4,312 test data points; see detailed count overview in Figure S2). TCRs in the training sets are distinct from those in the test sets but share the same epitope. The practical task can thus be viewed as finding patient-specific TCRs effective toward a well-known epitope (*e*.*g*., from SARS-COV2 spike protein). The IMMREP_2022 data splits were previously used to benchmark various ML methods and we report these results as baselines for TCRcube models. These baselines include peptide-specific sequence similarity models (tcrdist3^23^ and TCRbase^24^), shallow supervised ML models (DiffRBM^25^, SETE^26^, sonia^27^, tcrex^28^, TCRGRP^29^), supervised neural networks (netTCR^4^, TCRAI^30^, pMTnet^31^), and unsupervised language models (TCR-BERT^7^).

In Figure 2A, we show results for all methods considered, including all of our TCRcube models, aggregated across the 17 peptide-specific data splits. We observe that in general, ML methods highly outperform scoring function-based methods. Our TCRcube models are middle-of-the-pack when only raw amino acid identities are used as input (AAid) but improve considerably when residue positions are included (AAid + pos), becoming highly competitive with the best methods. When structure-aware AF2 latent representations are used, performance on average increases. Interestingly, AF2 structure module representations (AF2-strmod) slightly underperform evoformer ones (AF2-evoformer). The difference is not statistically significant, but might hint at the richer evoformer representations being more informative of TCR specificity. Moreover, more abstract representations derived from ESM protein language models lead to the best performance overall on the peptide-specific task, yielding a state-of-the-art TCRcube model. Notably, it is beneficial to extract ESM representations of the interface residues in the context of the full-length sequence of the TCR-pMHC complex (ESM2-full) instead of only using the relevant sequence fragments in isolation (ESM2-isol). In both instances, the same set of residues are being provided as input, but in the former case additional contextual information is encoded in the ESM embeddings. We note that, somewhat surprisingly, using the full pMHC and TCR sequences as input decreases performance (see Figure S3 for ablation experiments). Collectively, these results indicate clear gains using ML models over structure-based scoring functions for peptide-specific modeling.

**Figure 2.**
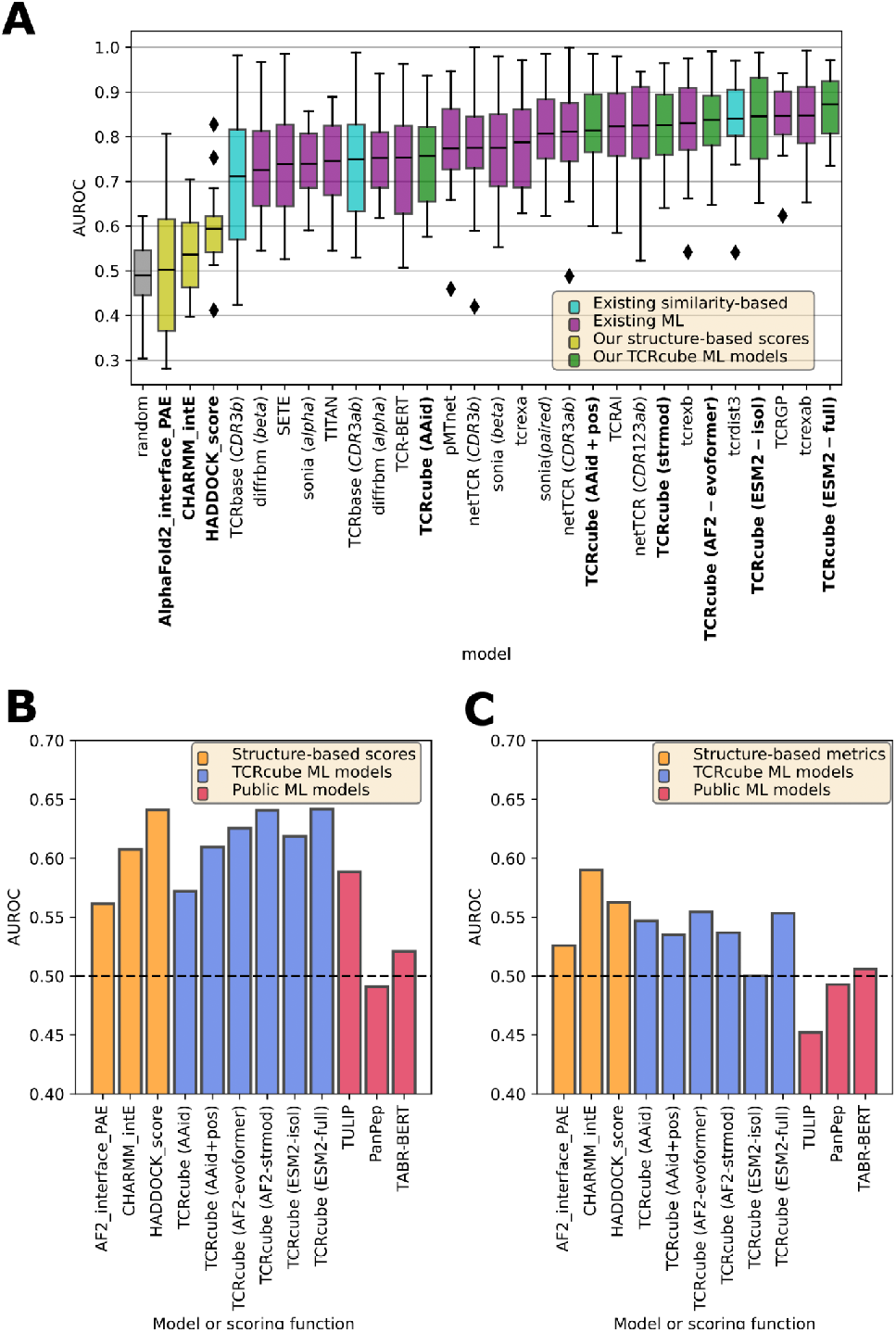
(**A**) Performance as measured by Area Under the Receiver-Operator Characteristic (AUROC) of scoring functions, peptide-specific ML models, and variants of TCRcube on the IMMREP_2022 test set. Each box and whiskers represents the aggregated results of 17 peptide-specific test sets. Lines correspond to mean values, while whiskers extend to the farthest data point lying within 1.5x of the inter-quartile range. (**B-C**) AUROCs of scoring functions, pan-peptide ML models, and variants of TCRcube on a (B) publicly available pan-peptide test set and a (**C**) proprietary neoantigen test set. Names of existing ML models are parenthesized with the following indicators of the input TCR representation: ‘alpha/cdr3a’ - CDR3*α*, ‘beta/cdr3b’ - CDR3*β*, ‘ab’ - combined CDR3*α* and *β*, ‘cdr123ab’ - all CDR loops and both *α* and *β*. Similarly for TCRcube models: amino acid identity - AAid, amino acid identity with addition of positional encoding - AAid+pos, AF2 evoformer single representations - AF2-evoformer, AF2 structure module single representations - AF2-strmod, ESM2-3B embeddings for isolated fragments - ESM2-isol, ESM2 embeddings for full complexes - ESM2-full.

### Pan-peptide evaluation of out-of-distribution generalization

Given the large number (>10^15^) of potential epitope peptides presented by MHC-I molecules^32^ and the correspondingly large sizes of human TCR repertoires (>10^8^ potentially promiscuous TCRs)^2^, the utility of peptide-specific models is somewhat limited. To achieve full utility, TCR-pMHC models must generalize to unseen peptides. To test current methods in this more general, pan-peptide setting, we designed a peptide-based training/test split of publicly-available TCR-pMHC pairing data composed exclusively of full-length sequences of the TCR-pMHC interface (79,086 training and 1,207 test data points; see Table S2 for full counts). To prevent data leakage of AF2 training data from our test sets, we excluded all binding data with corresponding experimentally-determined TCR-pMHC complexes in the Protein Data Bank. Independently, we also sought to assess methods in a completely blind fashion on a non-public data set. For this purpose we used a proprietary Genentech/Adaptive Biotechnologies data set on cancer neoantigens as a zero-shot test case (1,446 data points; see Table S3 for full counts). Note that public data comprises predominantly viral antigen complexes (Figure S4), enabling the proprietary test set to assess models in a novel cancer-focused regime.

As in the case of peptide-specific models, we predicted structures corresponding to all complexes in both training and test sets using the same evaluation pipeline as in the previous section (Figure 1). For this task, we restricted our attention to structure-based scoring functions (same set as before) and to pan-peptide ML models (Figure 2B, C - orange bars). As baselines, the ML methods of the previous section are not appropriate as they are designed in a peptide-specific manner. Instead, we selected three recent generalist ML models that reported promising performance on novel peptide generalization. Two models (TULIP^8^ and TABR-BERT^6^) are Transformer-based while the third (PanPep^5^) uses metalearning. For all three, we used checkpoints from their respective public repositories for making predictions (Figure 2B, C, magenta bars). Finally, we trained our TCRcube models using the new training set, which excludes all public test set peptides and proprietary peptides (Figure 2B, C, blue bars).

On pan-peptide test set based on public data (Figure 2B), we observe that structure-based scoring functions exhibit comparable performance to the peptide-specific setting This is unsurprising given the fact that they were not trained on either type of task and thus have uniform (albeit limited) predictive performance across the board. However, when these scoring functions are assessed on interactions involving cancer neoantigens (Figure 2C), we observe a noticeable drop in performance, suggesting a potential bias toward viral epitopes on which data is widely available.

The situation is markedly different for TCRcube and other ML models. The best TCRcube models, which in the peptide-specific setting substantially outperformed scoring functions, drop in performance to match the best scoring functions but no longer outperform them. It remains the case that ESM representations of the TCR-pMHC interface extracted in isolation (ESM2-isol) underperform their full-length complex counterparts (ESM2-full). However, contrary to peptide-specific models, AF2 structure module representations (AF2-strmod) slightly outperform AF2 evoformer representations (AF2-evoformer), although we do not have sufficient power to ascertain that this is statistically significant. Furthermore, in the specific case of cancer neoantigen prediction, the advantage of using representations derived from protein foundation models is diminished, with raw amino acid inputs being comparably performant, although all TCRcube variants do very poorly on this task. In fact, the predictive power of all approaches is low on the most challenging zero-shot task.

Turning to the three existing generalist ML models, we find that all of them exhibit low accuracies on the public test set. This is despite the fact that the training sets of all three models partially overlap with our public test set (23%, 21%, and 10% of test set epitopes were seen by TULIP, PanPep, and TABR-BERT, respectively). Thus, in contrast to our TCRcube models, this task is not zero-shot for the existing models. Yet only TULIP shows better than random performance on the public test set (Figure 2B). When the proprietary neoantigen test set is used, which is a zero-shot task for all three models, all fail to exceed even random performance and in fact do worse than random in some instances (Figure 2C).

These observations are inconsistent with previously reported results, which indicated that existing generalist models are capable of out-of-distribution generalization^5,6^. Here it is worth considering the training processes of the generalist models, all of which employed different strategies for simulating negative data points than the one we use for TCRcube models. Our approach relies on generating negative data points by mismatching TCRs to pMHCs within each data set separately to avoid contamination. PanPep and TABR-BERT, on the other hand, use TCR sequences from healthy patients as negatives, with the same pool of negatives for both training and test sets. We believe this may result in a form of data leakage and thus inflated performance, by allowing the ‘negative space’, common between training and test sets, to be used as the basis of discrimination by these models instead of the positive space. In other words, the models learn what does not bind as opposed to what does bind, and ultimately fail to generalize when presented with a genuinely novel test set. TULIP is the exception, as it is trained using an encoder-decoder scheme that relies exclusively on positive training data. Nevertheless, TULIP had the largest overlap between its training set and our test set, which may explain its comparatively inflated performance on the public test set.

### Performance of ML models as a function of peptide sequence distance

We sought to better characterize the degradation in predictive accuracy when transitioning between peptide-specific and pan-peptide models. So far our assessments have been binary, focusing exclusively on either in- or out-of-distribution generalization. We next assessed the performance of pan-peptide models as a function of the Levenshtein distance (Figure S5) between test set and training set epitopes on the public test set. As we needed fine-grained control over the training set in this evaluation, we focused exclusively on our TCRcube models. Unsurprisingly, we found that model performance decreases with increasing epitope distance when the training set is held fixed while the test set is stratified (Figure 3A). This behavior was consistent irrespective of the input representation used for a TCRcube model. We next held the test set fixed and modified the training set by gradually removing epitopes closest in sequence to the test set. This approach had the effect of shrinking the amount of data available for the model to train on. Here, performance dropped rapidly as well (Figure 3B). However, models that used protein foundation-based input representations exhibited a much more gradual decrease in performance, suggesting that they are indeed better able to generalize, at least to nearby regions of sequence space. Finally, we held the test set fixed and modified the training set by removing the most distant epitopes from the test set first (Figure 3C). This served the purpose of controlling for training set size reduction. Despite a precipitous drop in training set sizes, more dramatic than the experiment of Figure 3B, model performance was largely unaffected by the initial set of cuts, especially when using protein foundation models. Eventually, when training set size decreased by an order of magnitude, model performances dropped. Evidently, the closer the task is to the peptide-specific case (*e*.*g*., when the closest epitope in the test set is only one point mutation away), the better a model performs.

**Figure 3.**
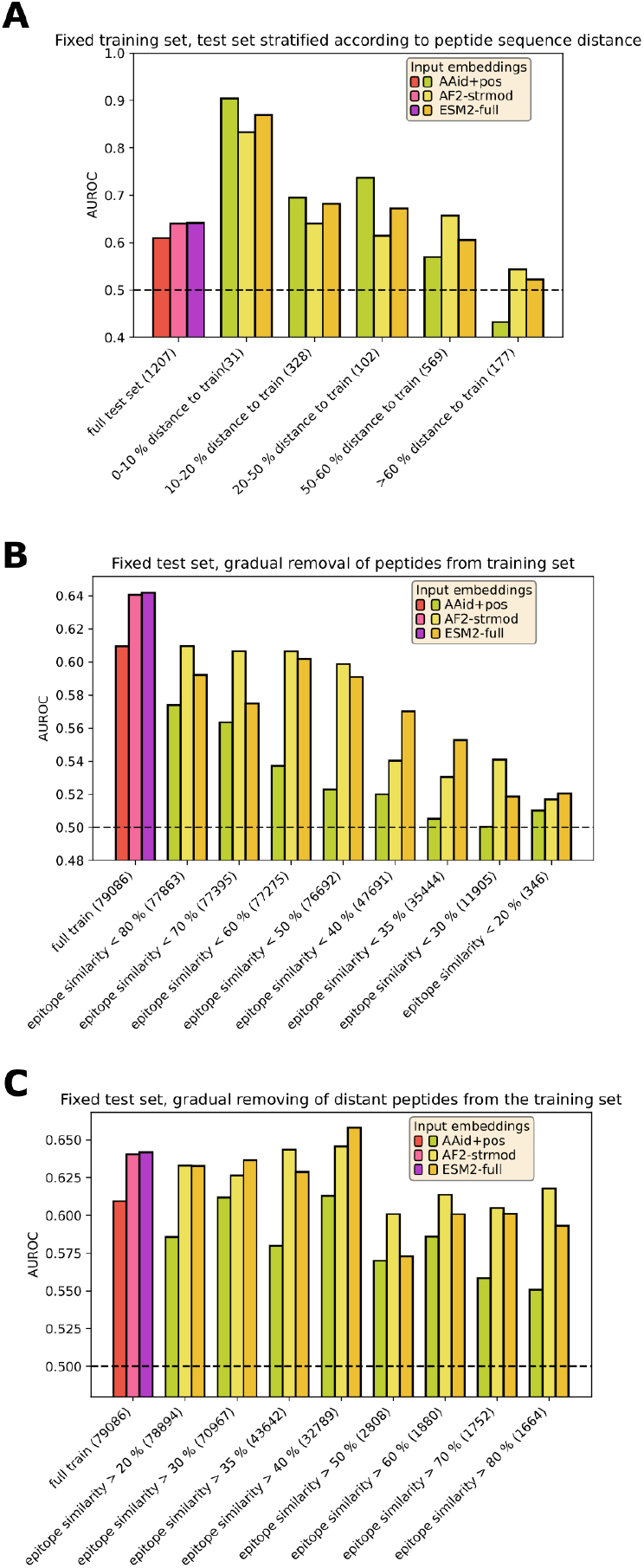
(**A**) AUROCs of TCRcube models assessed on test set subsets stratified by their epitopes’ Levenshtein distance to training set epitopes. (**B-C**) AUROCs of TCRcube models whose training sets have been altered by progressively removing epitopes (B) closest to or (C) furthest from the (full, original) test set.

### Strong TCR motifs lead to performant peptide-specific models

To better understand the basis of performant peptide-specific models, we searched for associations between model accuracy and aspects of the modeling task, hypothesizing that TCR sequence determinants may influence model behavior. To test this, we examined sequence similarity among TCRs, focusing on CDR3 loops in the peptide-specific IMMREP_2022 data sets. We aligned CDR3 sequences using the IMGT numbering scheme in ANARCI^33^ and constructed position-specific scoring matrices (PSSMs) of epitope-binding TCRs normalized relative to a background of non-binding TCRs. Strong sequence motifs in either CDR3*α* or CDR3*β* PSSMs were present in 4 out of 17 peptide-specific data sets. We show two examples in Figure 4A; for others see Figure S6 and S7. In those 4 data sets, the dominant sequence motif is present in the TCRs of 29% - 54% of the respective positive data points (Figure S8). When we compare performance of the 4 models containing a sequence motif with the remaining 13 without a clear motif, we find that TCR-motif-containing peptide-specific models are on the whole more performant than the motif-free ones, regardless of whether AF2- or ESM2 embeddings are used as the input representation (Figure 4B).

**Figure 4.**
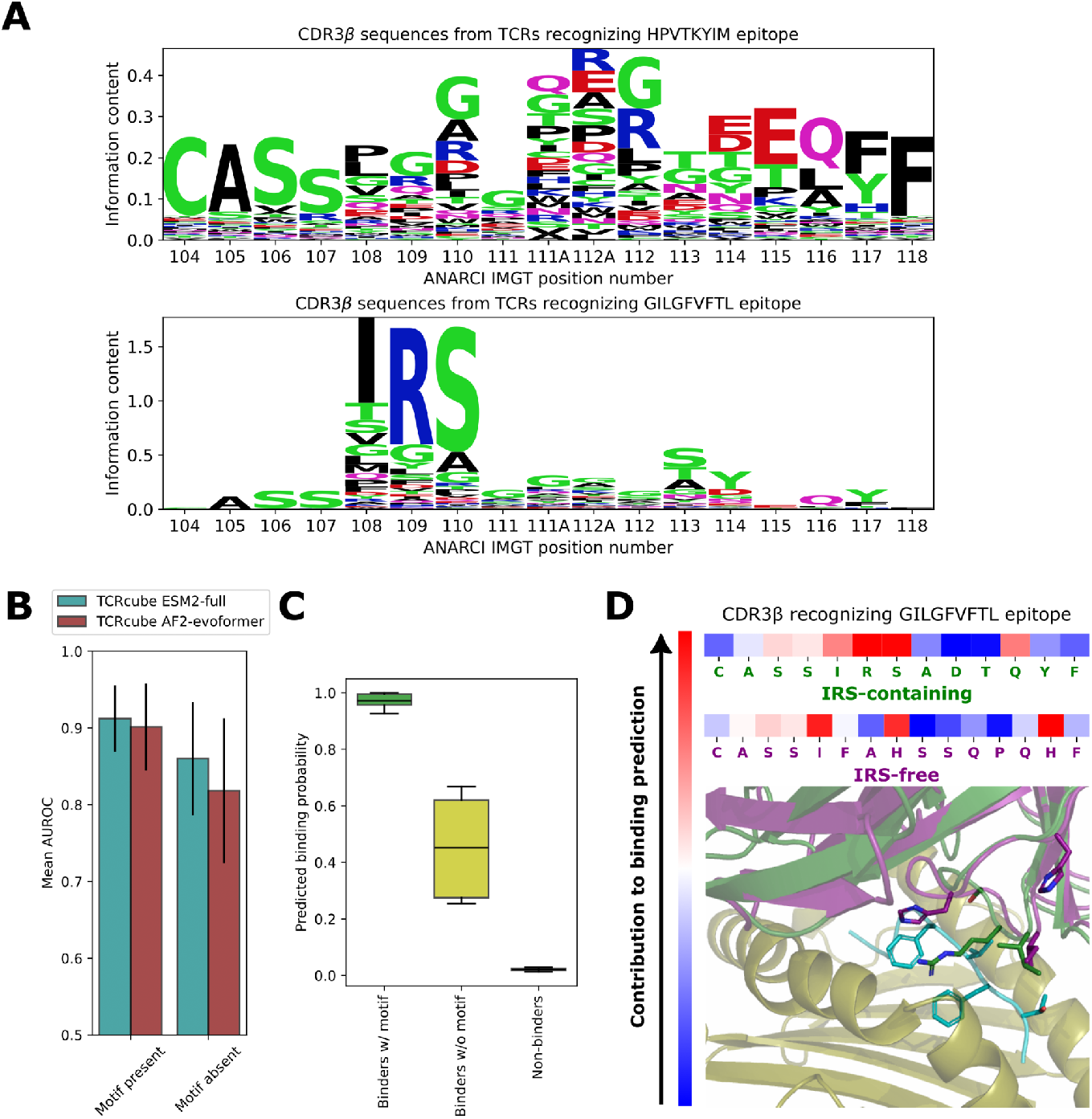
(**A**) Motifs in CDR3*β* sequences of TCRs in two peptide-specific data sets displayed as differential sequence logos (PSSMs of epitope-binding TCRs normalized relative to non-binding TCRs). (**B**) Mean AUROC of peptide-specific models stratified by the presence or absence of a dominant sequence motif. (**C**) Dependence of the predicted binding probability by the TCRcube ESM2-full model on the presence or absence of TCR motifs within 4 peptide-specific data sets (negative data points added as control). (**D**) Per-residue energy contributions from CDR3*β* residues of two distinct TCRs (top rows) with high predicted binding probabilities to GILGFVFTL epitope. First TCR (top row) contains the ‘IRS’ motif in CDR3*β* while the second (bottom row) does not. Contributions are based on the TCRcube AF2-evoformer model. AF2-predicted structures of both TCR-pMHC complexes are depicted below, with key interacting peptide (cyan) and TCR CDR3*β* (green or purple) residues shown in licorice representations.

We next assessed whether models rely on TCR motifs when making peptide-specific predictions. We compared TCR-pMHC binding probabilities within the 4 motif-containing models and found that models make more confident predictions for motif-containing TCRs (Figure 4C). Nevertheless, peptide-specific models are generally able to differentiate between positive and negative interactions, albeit with a smaller margin. When strong motifs are absent, model performance is weakly correlated with sequence similarity between training and test TCR repertoires (Figure S9).

To provide a specific example, we examined the contributions of individual CDR3 residues to the predicted binding probabilities of peptide GILGFVFTL, which comprises the single largest peptide-specific data set (4,094 data points) and has a clearly defined CDR3*β* ‘IRS’ motif (Figure 4A). When the ‘IRS’ motif is present, the model uses this information to identify the binding TCR, since the largest contribution to the overall binding energy prediction comes from the terms associated with the motif (Figure 4D). Looking at the structure prediction, it can be seen that the arginine in the ‘IRS’ motif interacts with the phenylalanines in the GILGFVFTL peptide, presumably through cation-*π* interactions^34^. When the motif is absent, however, the model finds alternative mechanisms of recognition: two histidines at different positions substitute for the arginine. We thus note that while dominant sequence motifs help performance, ML models are learning more advanced, and mechanistically plausible, interactions as well.

### Leveraging pan-peptide models to bootstrap peptide-specific learning

While pan-peptide models are generally less performant than their peptide-specific counterparts, their predictions are non-random and may capture general aspects of TCR-pMHC binding (Figure 2). We reasoned that this general knowledge may be leveraged in combination with peptide-specific training to achieve better performance. To this end, we retrained the peptide-specific HPVTKYIM model using pan-peptide model weights as a starting point, in lieu of randomly initialized weights. The HPVTKYIM peptide is absent from our pan-peptide training set, precluding data leakage. We observe a clear boost in accuracy for the majority of TCRcube model variants (Figure 5A). The HPVTKYIM peptide does not contain any strong TCR motifs, which suggests that it may be reliant on universal patterns of TCR-pMHC interactions, consistent with the observed boost. As a negative control, we repeat the same retraining process but with the previously discussed GILGFVFTL peptide, which does have a clear TCR motif. Since GILGFVFTL is included in the original pan-peptide training set, we remove all instances of it from the training set and train new pan-peptide models, which we use as initialization for training a GILGFVFTL-specific model. In contrast to HPVTKYIM, we see no performance boost from this approach (Figure S10), suggesting that motif-dependency precludes utilization of general TCR-pMHC knowledge in pan-peptide models.

**Figure 5.**
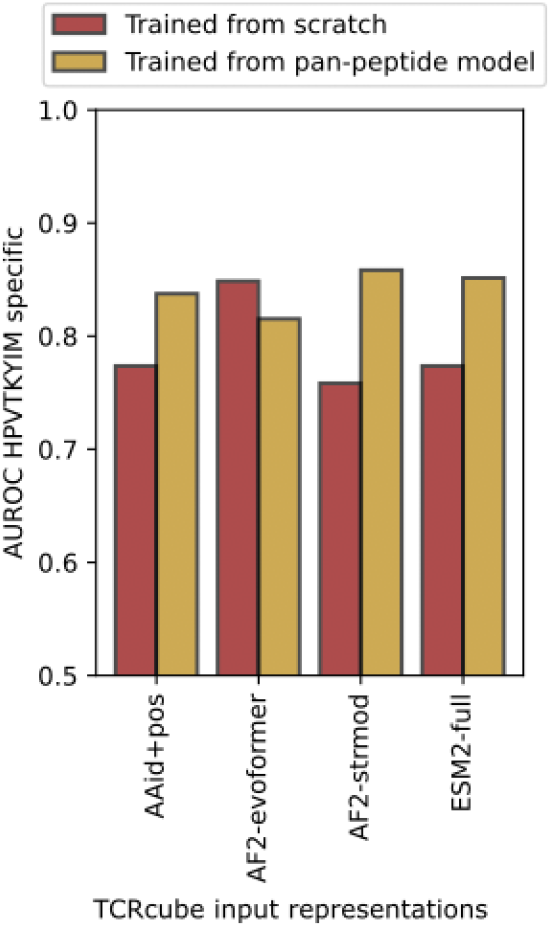
Impact of using pre-trained pan-peptide models as initialization for training peptide-specific variants of TCRcube models for the HPVTKYIM epitope.

## Discussion

Recent developments in ML-based TCR-pMHC modeling have suggested significant advances in our ability to identify TCRs specific to a given pMHC within an immune repertoire. In this work, we set out to investigate the extent to which ML models generalize to unseen epitopes relative to structure-based scoring functions. Apart from assessing existing models, we developed several variants of a relatively simple ML architecture and investigated its ability to leverage latent information encoded in large protein foundation models, specifically AlphaFold2 and ESM2, to predict TCR-pMHC specificity.

Despite substantial differences in their underlying architecture and training data, latent representations from both AlphaFold2 and ESM2 substantially improve the accuracy of our TCRcube models relative to simple representations based only on amino-acid identities. The resulting models achieved state-of-the-art performance on both peptide-specific and pan-peptide modeling, with high absolute accuracy on the former and low absolute accuracy on the latter. This is unlike classical, structure-based scoring functions, which show comparable (poor) performance on both tasks.

These results suggest that ML models learn peptide-specific aspects of TCR-pMHC interactions well but struggle to extract general principles of binding. Given the comparable performance of TCRcube and scoring function-based methods on pan-peptide tasks, we are led to conclude that the utility of protein foundation models to interaction prediction is comparable to that of physically-grounded scoring functions, at least for tasks in which residue-residue co-evolution, commonly used in structure and complex prediction, is minimal or absent. On the other hand, the success of peptide-specific models appears rooted in learning specific TCR sequence motifs that underpin peptide recognition, perhaps in combination with more general patterns of interaction, and the latter can be aided by pretraining on pan-peptide tasks. The performance of peptide-specific TCRcube models can thus be explained by a combination of general protein knowledge extracted from foundation models (AF2 or ESM2), general TCR-pMHC interaction knowledge from pan-peptide models, and peptide-specific knowledge to a certain extend dependent on sequence motifs.

The difficulty that pan-peptide models exhibit in generalizing to novel epitopes suggests that additional data, and perhaps additional types of data, extending beyond sequences and static structures might be necessary to capture the underlying physical basis of TCR-pMHC interactions. Indeed, it has been shown that pMHC recognition by TCRs is dynamic^35,36^, with activation dependent on unbinding kinetics rather than affinity^37–39^, with the catch bond^38^ or kinetic proofreading^40^ mechanisms being leading hypotheses. Hence, scoring function-derived energies calculated based on static structures, whether experimental or predicted by AlphaFold2, may fundamentally not be predictive of activation, necessitating approaches that capture differences in dynamical behavior between the interfaces of productive and non-productive complexes. Alas, TCR-pMHC data with quantified thermodynamic and kinetics parameters remains exceedingly limited.

Furthermore, the majority of publicly available data employs a specific TCR discovery strategy^41^ based on multimer binding assays that has been in use for several decades. Consequently, existing data may well be biased toward ‘stronger binders’,^42^ constraining the range of functionally relevant interaction patterns that can be learned by ML models, as it has been shown that weak binders can also activate T-cells and strong binders do not always do so.^38^ Moreover, even if T-cells are screened for activation in addition to binding, the dependency of T-cell activation on pMHC concentration^43^ adds further noise to the data, since improper calibration of experimental pMHC concentrations can lead to false negatives. Capturing the broad determinants of TCR specificity thus likely requires generation of a data set that simultaneously reflects both binding and activation, ideally at varying concentrations of TCRs and pMHC molecules. This approach would also potentially accommodate empirical evidence supporting the kinetic proofreading hypothesis.

It is yet not clear what architecture will be ideal for modeling the kinetic aspects of TCR-pMHC interactions. Recognizing that the nature of the TCR-pMHC interfaces is dynamic^35^, integration of MD simulations into the pipeline^46^ or leveraging generative ML models^47^ able to reproduce the conformational ensembles from MD simulations hold some promise. However, the performance of those models are still on their way to reach the level of accuracy that AlphaFold2 reached for the single-structure prediction task.

## Methods

### Data

#### Public data

For peptide-specific evaluation, we used the IMMREP_2022^17^ TCR specificity benchmark set (downloaded from thre respective GitHub repository https://github.com/viragbioinfo/IMMREP_2022_TCRSpecificity). This data set contains ready-to-use training and test sets for 17 peptides curated from public sources with full information about TCRs needed for structural modeling (paired CDR3*α* and CDR3*β* sequences and full gene annotation). For full pMHC modeling, we had to search for the respective MHC allele in the VDJdb database^48^. Furthermore, the repository contains results of various published methods. We use those unmodified previous results as a baseline for evaluation of herein-presented analyses and ML models.

For construction of the pan-peptide data set for general TCR specificity, we used data from public sources curated in VDJdb^48^ (downloaded 04/10/2023), Mc-PAS-TCR^49^ (downloaded 03/15/2023), and IEDB^50^ (downloaded 03/06/2023) as well as data coming from the 10x Genomics public resource. We restricted our data to the complexes corresponding to human MHC-I alleles, and selected specifically the data points with paired CDR3*α* and CDR3β information and fully-annotated V and J genes (TRAV, TRAJ, TRBV and TRBJ). Full TCR information is necessary in order to reconstruct a full sequence of the variable domain for structural modeling. In order to account for potential bias from AlphaFold2 training, we excluded all data points present in the PDB. We selected data points corresponding to 62 epitope peptides with a maximum of 50 known TCR interactors for the test set for the general TCR specificity task (205 interactions). The remainder of the positive data points corresponding to 732 epitopes (13,181 interactions) were kept for the training set for herein-presented ML models. For both training and testing, non-productive (negative) pairings are required. Those data points are, however, scarce in the literature and negative pairs corresponding to just a handful of epitope peptides are available. To ensure uniform coverage of the space, we generated negative data points within each training / test split of the data set by assigning different peptide-MHC pairs to each unique TCR. We generated 5 negative data points per one positive data point within each of the sets by assigning different epitopes to each unique TCR. After filtering out potential duplicates, this resulted in 1,207 test and 79,086 training data points. For the performance analyses, we calculated minimal Levenstein distance (normalized by peptide length) between the training and test peptides (*i*.*e*., determined which peptide from the training set is closest to the given test set peptide). We then used this information to evaluate performance on the test set using distance-based subsets. We also divided the training set in a similar way and used either sequence-proximal or sequence-distant portions of the training set for model training.

#### Lung cancer-derived internal data

The second (internal to Genentech/Adaptive Biotechnologies) pan-peptide test set is based on experimental identification of TCR-neoantigen pairs in 30 non-small cell lung cancer (NSCLC) patients from the previously-described Lung Cancer Mutation Consortium 3 (LCMC3) trial (NCT02927301)^51^. Briefly, this was a phase II, open-label, single-arm study of neoadjuvant atezolizumab followed by optional adjuvant atezolizumab and/or chemo- or radiotherapy. Eligibility criteria included age ≥18 years, pathologically confirmed stage IB-IIIA or select IIIB NSCLC, confirmation that the primary tumor and involved lymph nodes were technically resectable, ECOG PS 0-1, and signed informed consent to participate in the study. Before treatment, a primary tumor core biopsy and blood samples were taken for biomarker analyses.

#### Neoantigen prediction in 30 non-small cell lung cancer patients

Neoantigens were predicted from tumor/normal whole exome sequencing and tumor RNAseq. In brief, WES and RNAseq data were aligned using GSNAP^52^ (v2013-11-01), with WES realigned using GATK IndelRealigner^53^ (v3.8-0-ge9d806836) and mutations were called using both lofreq2 (v2.1.2) and strelka1 (v2.0.17.strelka1). Evidence for RNA expression of mutant alleles was assessed using VariantTools^54^ (v1.28.1). Mutations with support from > 2 RNAseq reads were retained for further characterization. Coding mutations were identified and translated *in silico* using VEP^55^ (release 90) with the Downstream plugin. Mutant sequences along with 12mer wild-type flanking sequences on either side were extracted as putative neoantigens. Patient class I HLA alleles were identified using HLA-HD^56^ (v1.2.0.1), and the constituent 8-11mer neoepitopes of the putative neoantigens were scored for MHC presentation using NetMHCpan4.0^57^. Up to 80 neoantigens were selected for experimental TCR identification per patient by ranking on the sum of ranks according to NetMHCpan scores and mutation VAFs.

#### Neoantigen-specific TCR identification using MIRA

TCR-neoantigen pairs were identified using Multiplex Identification of T-cell Receptor Antigen Specificity (MIRA) – a multiplexed assay in which subsets of patient T-cells are stimulated with unique mixtures of pooled antigens. Antigen responsive T-cells are then isolated via flow cytometry using markers of T-cell activation and their TCRB locus sequenced^58^. Antigen pairing is achieved using a nonparametric Bayesian model described previously^59^ that identifies TCRβ sequences that coincide with the occupancy of particular antigens^60,61^. MIRA was performed using PBMCs derived from 30 NSCLC patients in the LCMC3 trial. In brief, patient-derived PBMCs were stimulated with CD3:CD28 for polyclonal expansion of peripheral T-cells. Patient T-cells were then transfected with neoantigen-encoding RNA constructs in a 9-choose-5 format, such that each neoantigen corresponded to a unique 5-well address and 9 total antigen exposures. Responsive T-cells were then isolated using FACS based on CD137 expression, followed by immunosequencing of the TCRB locus. Coincidences of TCRβ sequences with neoantigen addresses were used to establish preliminary TCR-antigen pairs. In parallel, TCR*αβ* pairs were established using pairSEQ^62^. TCR-antigen specificities were orthogonally verified by transiently transfecting Jurkat cells with RNA constructs encoding the putatively neoantigen-specific TCR*αβ* and co-incubating them with K562 cells transiently transfected with each of the individual patient-specific class I HLA alleles and neoantigen constructs. Responsiveness was assessed by Jurkat expression of CD69, and was compared to the wild-type counterpart of the mutant neoantigen. From this process, 241 data points with full TCR gene annotation and paired CDR3 sequences, and verified neoantigen specificity were selected, and 5 negatives per positive data point were generated as described above, totaling 1,446 data points. Since the MIRA assay was performed using transgenes encoding 21mer neoantigens, epitope-level resolution of TCR specificity was not available, and NetMHCpan-4.1^63^ was used to approximately determine the MHC-presented epitope peptide.

### AlphaFold2 predictions of TCR-pMHC complex structures

We used the TCRdock^20^ pipeline for AlphaFold2^19^ predictions of the structures of the interface domains (TCR and MHC variable domains near the interface) of all TCR-pMHC complexes in our test and training sets (both positive and negative data points). TCRdock constructs 4 patchwork templates based on public structures from the PDB. These are used alongside the query sequence as the sole input for AlphaFold2 (*i*.*e*., no MSA is required). Given the hypervariable nature of the regions at the TCR-pMHC interface, the relevance of MSAs for TCR-pMHC complexes is disputable. Contrary to the default setup, we ran only one prediction per data point. Otherwise, we use default settings, including *model2* parameters and three recycles. We also modified the TCRdock code to output evoformer single and structure module single latent representations (sequence_length x 384 tensors) in addition to the structure and confidence metrics (plDDT and PAE) for each of the complexes. Both latent representations are extracted after passing through all the evoformer and/or structure module blocks in the last recycle.

For the evaluation of the predictive performance of the AF2 metrics, we use predicted alignment error (PAE) at the TCR-pMHC interface. The interface is defined by CDR3 loops and peptide residues.

### Calculation of force field interaction energies

Each AlphaFold2 prediction of a TCR-pMHC complex structure was prepared for interaction energy calculation using vmd^64^ 1.9.3 and its modules. All steps were done using tcl scripts with default settings to allow processing at scale. We used the *psfgen* module with default settings to add hydrogens and force field topology (using CHARMM 36 force field^21^). We then added a 10Å water box around the protein complex using the *solvate* module. We subsequently used NAMD^65^ 3.0 to minimize this structure of the TCR-pMHC complex in 1,000 steps. The aim was to resolve potential clashes and allow the water molecules to move to a more realistic position. We then used the *interaction* keyword in NAMD to calculate the non-bonded (electrostatic and van der Waals) interaction energy between the pMHC and TCR part of the complex.

### Re-docking of TCR-pMHC complexes

We used HADDOCK^22^ 2.5 for re-docking TCR-pMHC complex structures. As an input, we take the pMHC and TCR parts of the AlphaFold2 prediction. We consider all residue pairs from the peptide and TCR that are within 4 Å in the AlphaFold2 structures as active for the HADDOCK protocol. We modify the standard protocol to increase computational feasibility at large scales: we decrease rigid body docking attempts to 300, number of structures for refinement to 20, number of trials for rigid body minimization to 3, number of MD steps to 50 for the rigid body part and 200 for the flexible part, number of for heating phase to 20 in solvated rigid body docking, sampling steps to 250, and cooling steps to 200. We take the structure with the best HADDOCK score as representative for each data point.

### Pre-calculation of ESM2 embeddings

We extracted ESM2 representations either for a separated sequence of CDR3 loops, peptide, and variable domain of MHC, or the full sequence of the TCR-pMHC complex (as used for AlphaFold2 predictions). In the case of a full-length sequence of the complex, we used 50-glycine-residue linkers to connect the individual proteins of the complex. Linker usage is a recommended strategy for multimer treatment proposed in the ESMfold^13^ code (a similar strategy has been used previously to predict multimeric complexes using the monomeric version of AlphaFold2^66^). ESM2^13^ representations were extracted using scripts provided in the ESM github repository. We used the per_tok representations of the last (36^th^) layer of the esm2_t36_3B_UR50D version of the ESM2 protein language model (each residue is represented by a 2,560 dimensional embedding). This ESM2 version has shown the best performance/speed ratio for the ESMfold^13^ model.

### TCRcube machine learning model architecture

We used pyTorch^67^ to implement the TCRcube architecture (Algorithm 1, schematically depicted in Figure 1). As the input, TCRcube takes the residue-level representations of the sequences of the epitope peptide and the variable part of MHC and CDR3*α* and CDR3*β* sequences of TCR. The representations used in this paper are either AlphaFold2 latent representations (evoformer or structure module single representations) or sequence-only-based representations in the form of ESM2 embeddings or simple amino acid identities (nn.Embedding module in pyTorch was used for both the identity as well as position in the sequence fragment). Any type of representation is first projected into the inner dimension (384). The model then works in two stacks corresponding to the TCRɑ and TCRβ components. The TCR representation is combined with the peptide and MHC representations through an outer product, which results in the following tensor: (TCR_seq x MHC_seq x peptide_seq x embedding_dim). This representation corresponds to a latent representation of the interactions on the peptide-MHC-TCR interface and should capture not only the peptide-TCR part, but also the peptide-induced MHC interactions. We then apply layer norm, ReLU activation, and a linear layer, which project the above tensor from embedding dimension to 1. Thus we get a set of peptide-MHC-TCR energy terms. We then calculate the sum of those terms and normalize it by the length of the peptide, MHC and CDR3 sequences in order to account for different sequence lengths across the data set.

#### Algorithm 1.

TCRcube model

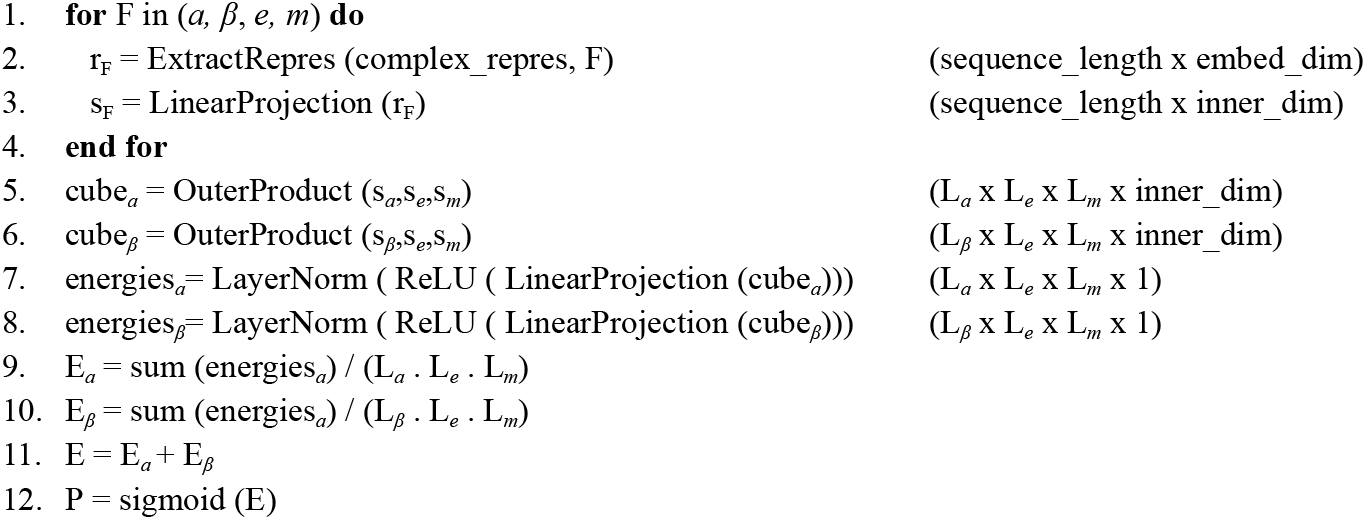

### TCRcube machine learning model training process

The models were trained as binary classifiers using binary cross entropy as a loss function. We upweighted positive data points to reflect the positive:negative ratio in the training data set. We used Adam optimizer with an initial learning rate of 10^−4^ and default parameters otherwise. We trained all models for 500 epochs to determine the point of maximal performance on the validation set and took this checkpoint for any further performance evaluation.

### Evaluation of the predictive power of ML models and structure-based scoring functions

Given the nature of available data, we evaluated all predictions using binary classification (discriminating between productive / non-productive TCR-pMHC interactions). With respect to how the results are reported in the field of TCR-pMHC ML models, we have chosen the area under the receiver operating curve (AUROC) as the reported metric. ROC curve represents the relation between true positive and false positive rates under various classification thresholds. The area under the ROC curve represents the mean performance of the classifier under all thresholds. It has been shown that AUROC is actually the best choice when evaluating a binary model in threshold-agnostic setting.^68^

### Predictions using public models

We downloaded the inference code and pre-trained model checkpoints of TULIP^8^, PanPep^5^ and TABR-BERT^6^ models from their respective github repositories (as referenced in the papers). We used all models in their zero-shot mode to make predictions on both of our test sets. Scores coming from each model were used to calculate AUROC.

### Analysis of sequence motifs in TCRs

We used the ANARCI tool^33^ to number CDR3 sequences according to the IMGT scheme and placed gaps at the positions without an assigned amino acid. We used the *logomaker* package in python to produce information content matrices (*to_type=‘information’*) from aligned sequences of positive data points while using the PSSM of negative TCRs as background. We then used the same package to visualize sequence logos, while we elided positions with less than a half non-gap entries. Furthermore, we analyzed the underlying information content matrices to detect sequence motifs in the CDR3 sequences. We considered every amino acid with information content > 0.3 as dominant. For a sequence motif to be detected, we have to detect at least two such highly occurring amino acids.

## Acknowledgements

The authors would like to thank Adaptive Biotechnologies for collaboration in generating the lung-cancer-patient data set and providing the MIRA assay protocols.

## Data and code availability

The code and weights of the TCRcube model, as well as the training and testing data sets assembled from public resources are available on github: https://github.com/aqlaboratory/TCRcube. The proprietary lung-cancer-patient (lcmc3) data is licensed from a third party and the author’s employers do not have the right to provide this data for publication.

