## Supplementary Information for "Predicting specificity of TCR-pMHC interactions using machine learning and biophysical models"

Martin Culka[1], Nicolas W. Lounsbury[2], William Thrift[2], Santrupti Nerli[2], Andrew Wallace[2], Gergő Nikolényi[1], Darya Orlova[2], Kiran Mukhyala[2] and Mohammed AlQuraishi[1]

[1] Department of Systems Biology, Columbia University, New York, New York, 10032, USA

[2] Genentech, South San Francisco, California, 94080, USA

### Supplementary tables

**Table S1**. Overview of published pMHC-TCR machine-learning models

| Name | Input | Training scheme | Model architecture | Reference |
|---|---|---|---|---|
| NetTCR 2.2 | CDR1,2,3 $\alpha,\beta$ and peptide sequences | Specific / panpeptide | CNN | (Jensen and Nielsen, 2024) |
| PanPep | CDR3$\beta$ and peptide sequences | Panpeptide (healthy negatives) | MLP, metalearning | (Gao et al., 2023) |
| TULIP | CDR3 $\alpha,\beta$ , peptide and MHC allele | Specific / panpeptide | Transformer | (Meynard-Piganeau et al., 2024) |
| TABR-BERT | CDR3$\beta$ and peptide sequences | Panpeptide (healthy negatives) | Transformer | (Zhang et al., 2024) |
| RACER-m | Crystal structure and peptide sequences | Panpeptide (TCR-restricted) | Force-field with ML-optimized parameters | (Wang et al., 2024) |
| TEIM | CDR3$\beta$ and peptide sequences | Panpeptide | CNN + autoencoder | (Peng et al., 2023) |
| EPIC-TRACE | CDR3 $\alpha,\beta$ and peptide sequences | Specific / panpeptide | CNN + attention | (Korpela et al., 2023) |
| ERGO-II | CDR3 $\alpha,\beta$ and peptide sequences, VD genes, MHC allele | Specific / panpeptide | LSTM / autoencoder | (Springer et al., 2021) |
| TITAN | CDR3$\beta$ and peptide sequences | Specific / panpeptide | Cross attention | (Weber et al., 2021) |
| DLpTCR | CDR3 $\alpha,\beta$ and peptide sequences | Penpeptide (not epitope zero-shot) | Ensemble model | (Xu et al., 2021) |
| tcrex | CDR3$\beta$ and peptide sequences, VD genes | Specific | Random forest | (Gielis et al., 2019) |
| TCRGP | CDR3 $\alpha,\beta$ and peptide sequences | Specific | Gaussian process | (Jokinen et al., 2021) |
| SONIA | CDR3 $\beta$ and peptide sequences | Specific | MLE | (Sethna et al., 2020) |
| DiffRBM | CDR3 $\beta$ and peptide sequences | Specific | Restricted Bolzmann Machines | (Bravi et al., 2023) |

| | | | | |
|---|---|---|---|---|
| TCRAI | CDR3 $\alpha,\beta$ + VD genes | Specific | CNN | (Zhang et al., 2021) |
| TAPIR | CDR3 $\alpha,\beta$ + VD genes, peptide, MHC allele | Panpeptide | CNN | (Fast et al., 2023) |
| SETE | CDR3 $\beta$ and peptide sequences | Specific | Gradient Boosting Decision Tree | (Tong et al., 2020) |
| pMTnet | CDR3 $\beta$ and peptide sequences | Panpeptide | Autoencoder | (Lu et al., 2021) |

**Table S2**. Overview of data used for construction of public pan-peptide data set

| | Train set | Test set |
|---|---|---|
| Data points | 79086 | 1207 |
| Epitopes | 732 | 62 |
| TCRs | 13077 | 174 |

**Table S3**. Overview of data in the proprietary test set

| | |
|---|---|
| Data points | 1446 |
| Epitopes | 74 |
| TCRs | 241 |

### Supplementary figures

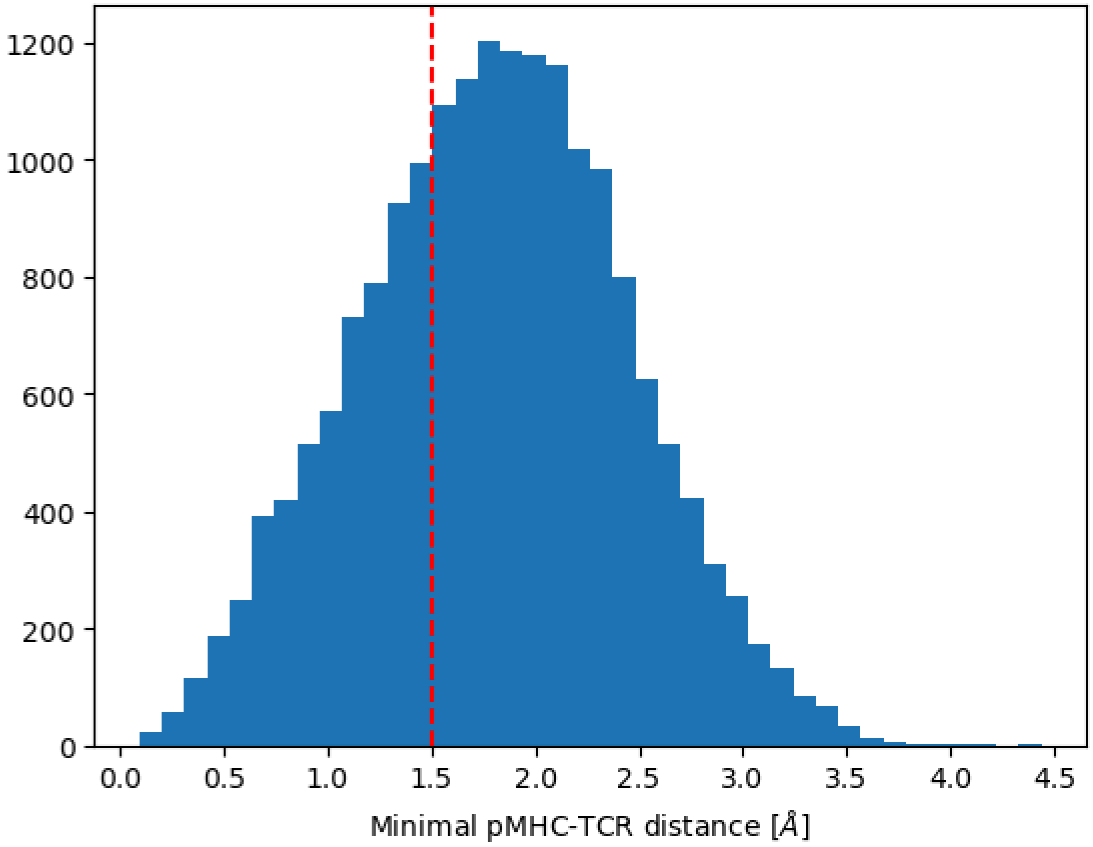


Figure S1. Minimal distance between pMHC and TCR atoms across the IMMREP_2022 data set. Red line represents a very conservative threshold for a clash (distance of heavy atoms < 1.5 Å).

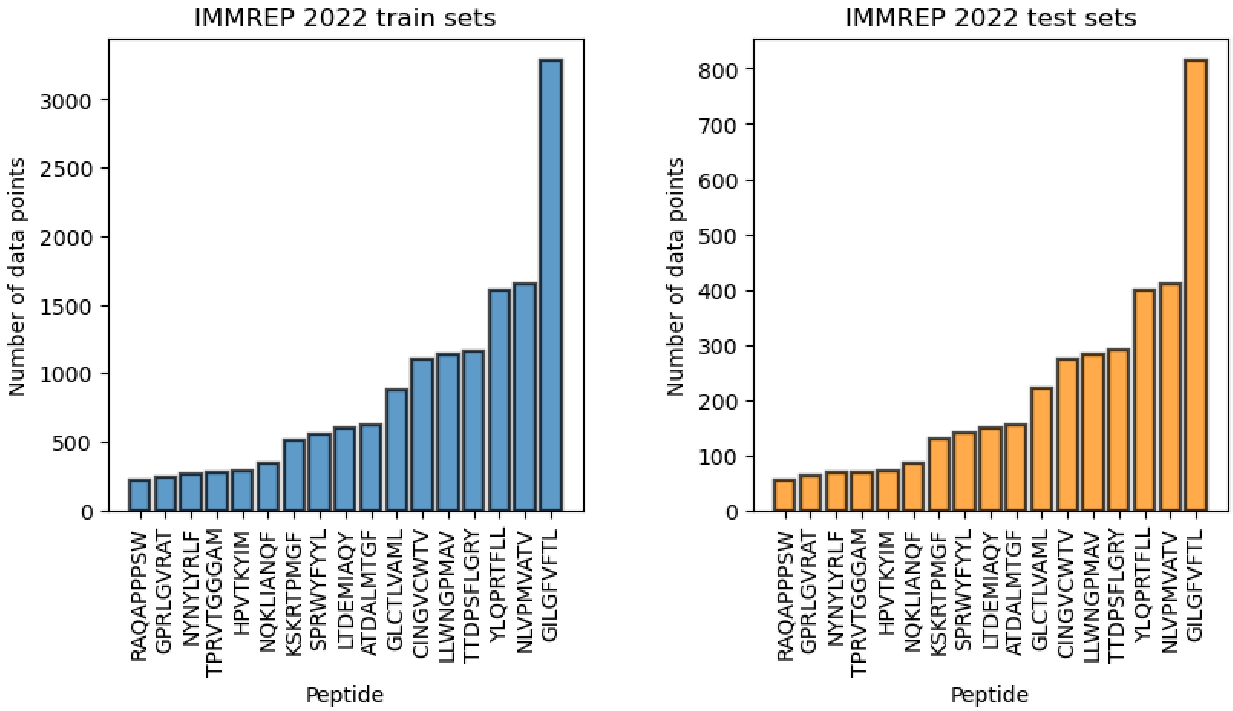


Figure S2. Overview of data point count in the peptide-specific train and test sets of IMMREP_2022.

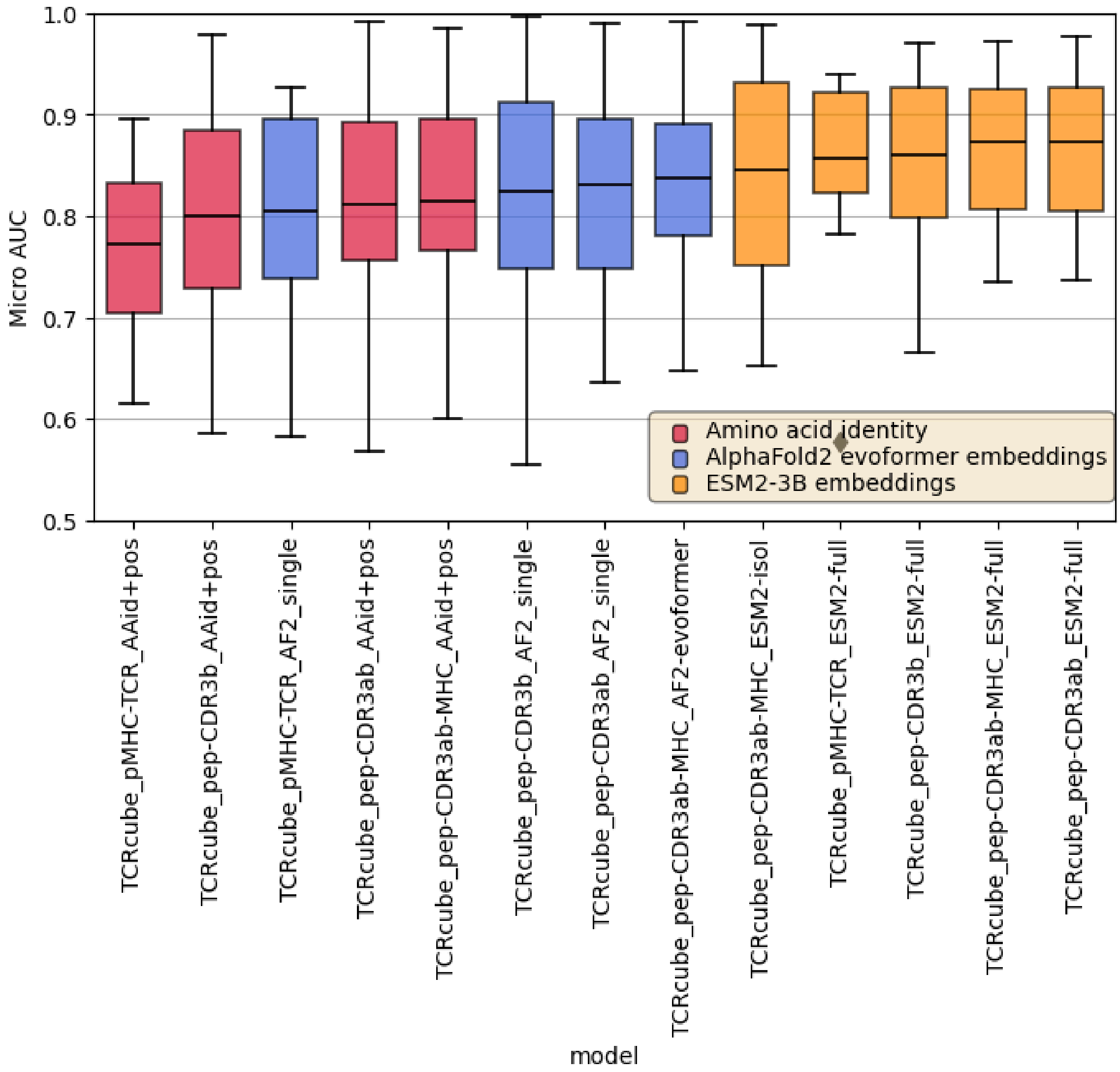


Figure S3. Ablation study on the effect of different types and extent of pMHC and TCR representations on the set of peptide-specific models trained on IMMREP_2022 data set.

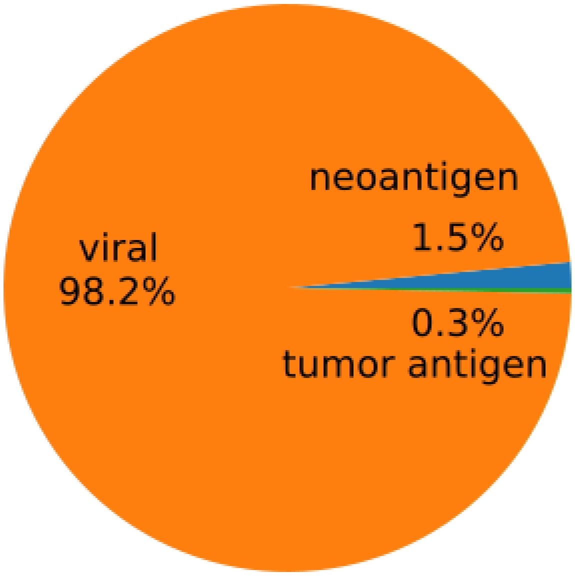


Figure S4. Share of categories of epitope peptides in our pan-peptide public data set (not all data points have origin labels).

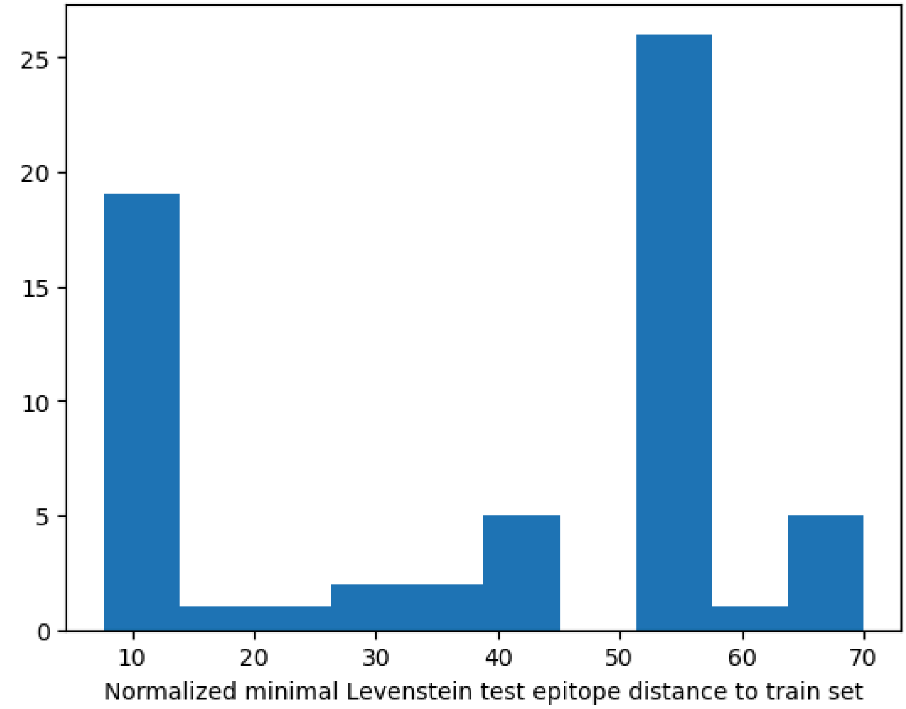


Figure S5. Histogram of minimal distances of test set epitopes to train set epitopes in the pan-peptide data set.

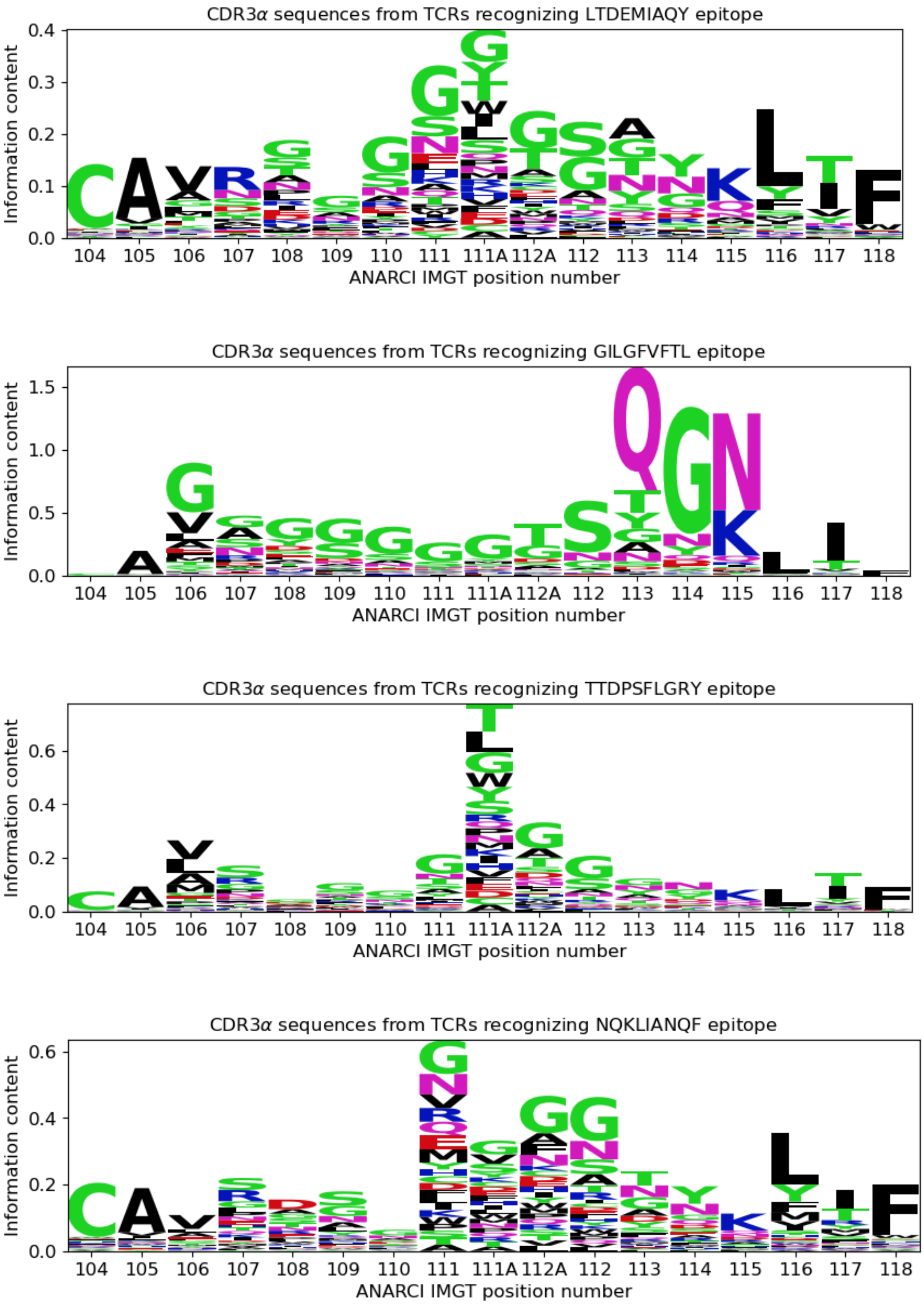


Figure S6. Differential sequence logos of peptide-specific TCRs from IMMREP_2022 data sets - CDR3$\alpha$

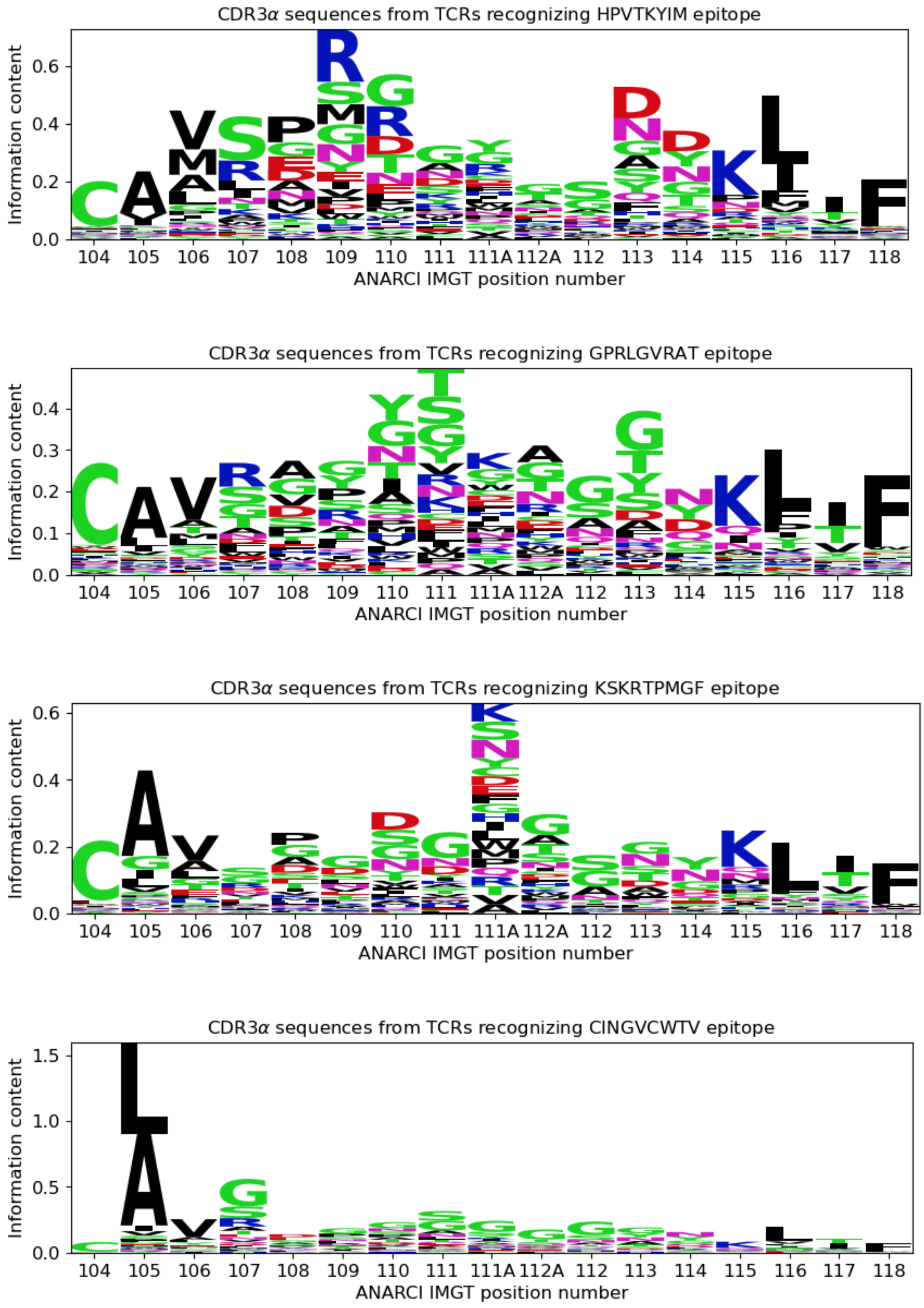


Figure S6. Differential sequence logos of peptide-specific TCRs from IMMREP_2022 data sets - CDR3$\alpha$ (continued)

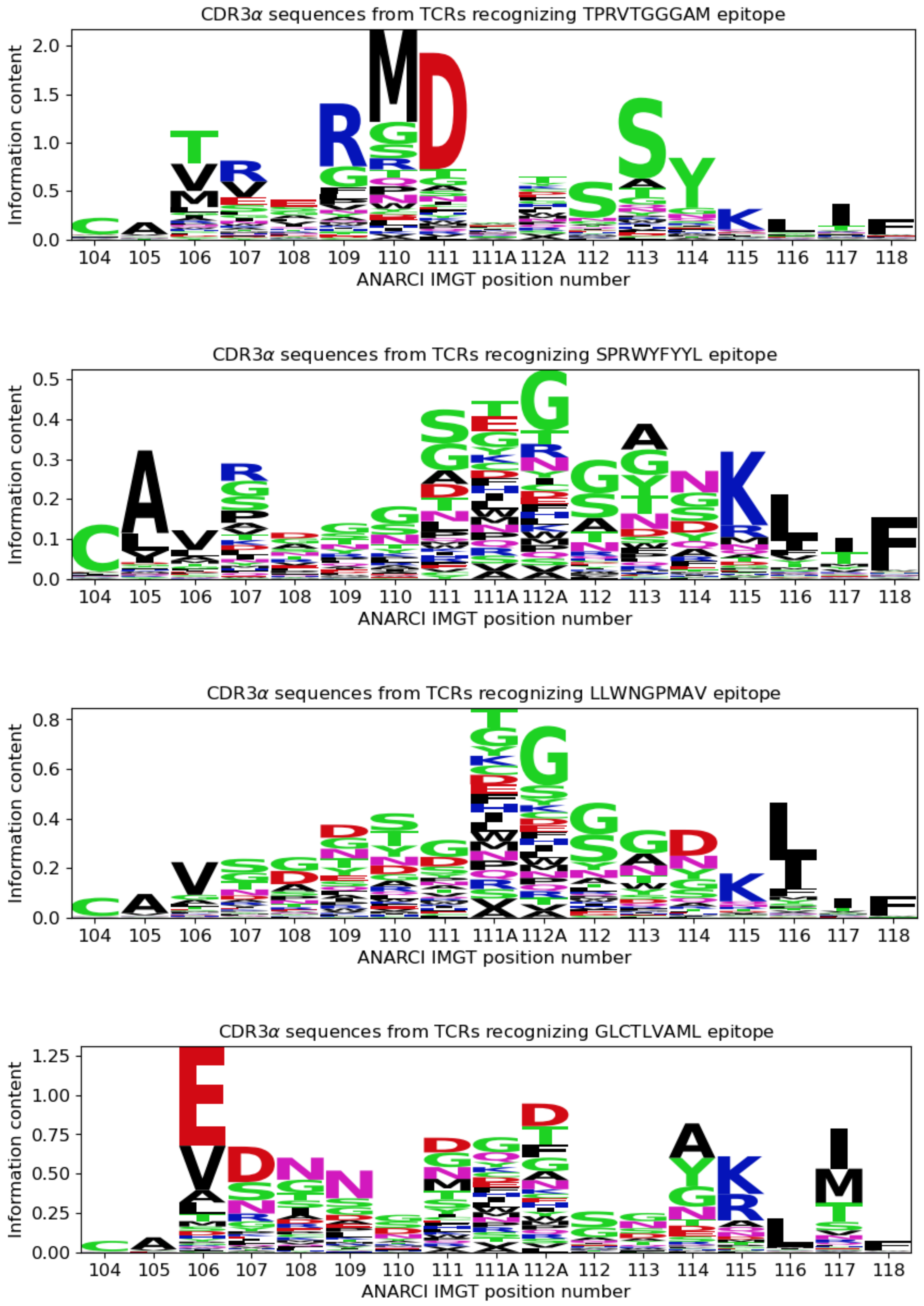


Figure S6. Differential sequence logos of peptide-specific TCRs from IMMREP_2022 data sets - CDR3α (continued)

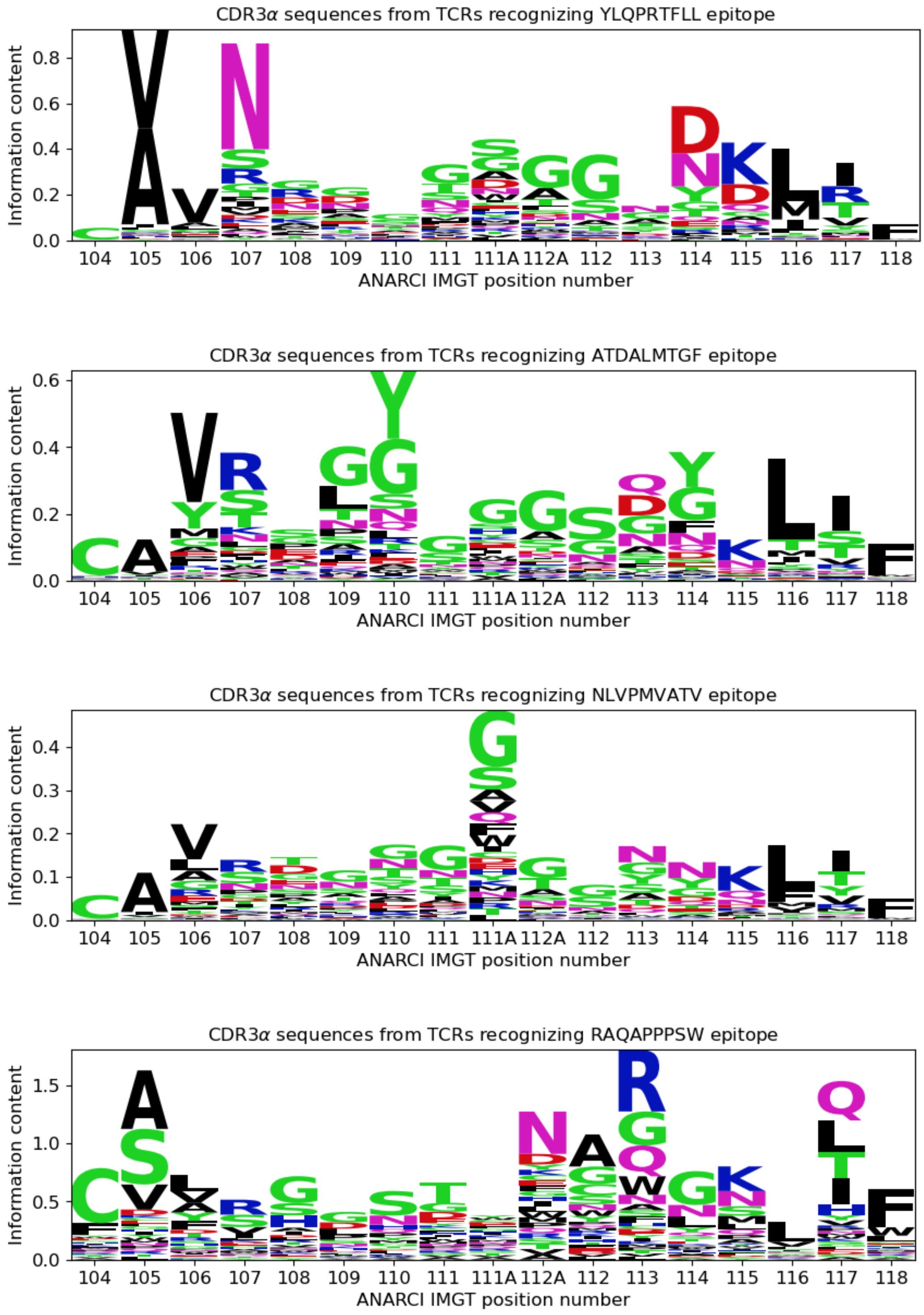


Figure S6. Differential sequence logos of peptide-specific TCRs from IMMREP_2022 data sets - CDR3$\alpha$ (continued)

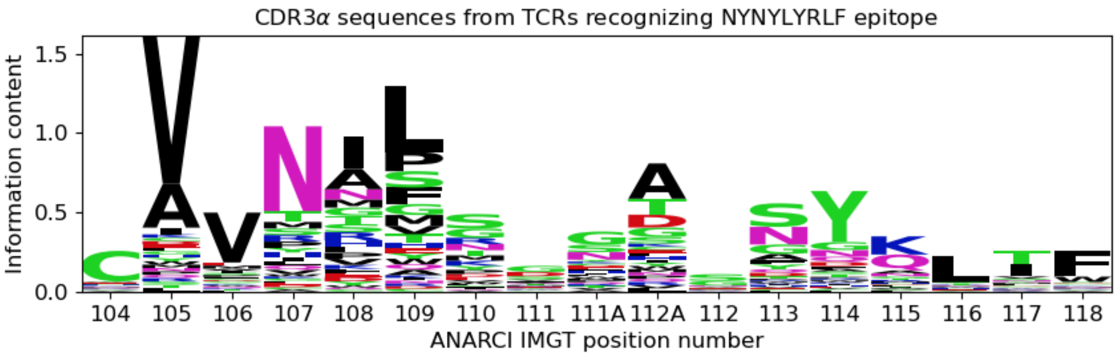


Figure S5. Differential sequence logos of peptide-specific TCRs from IMMREP_2022 data sets - CDR3$\alpha$ (continued)

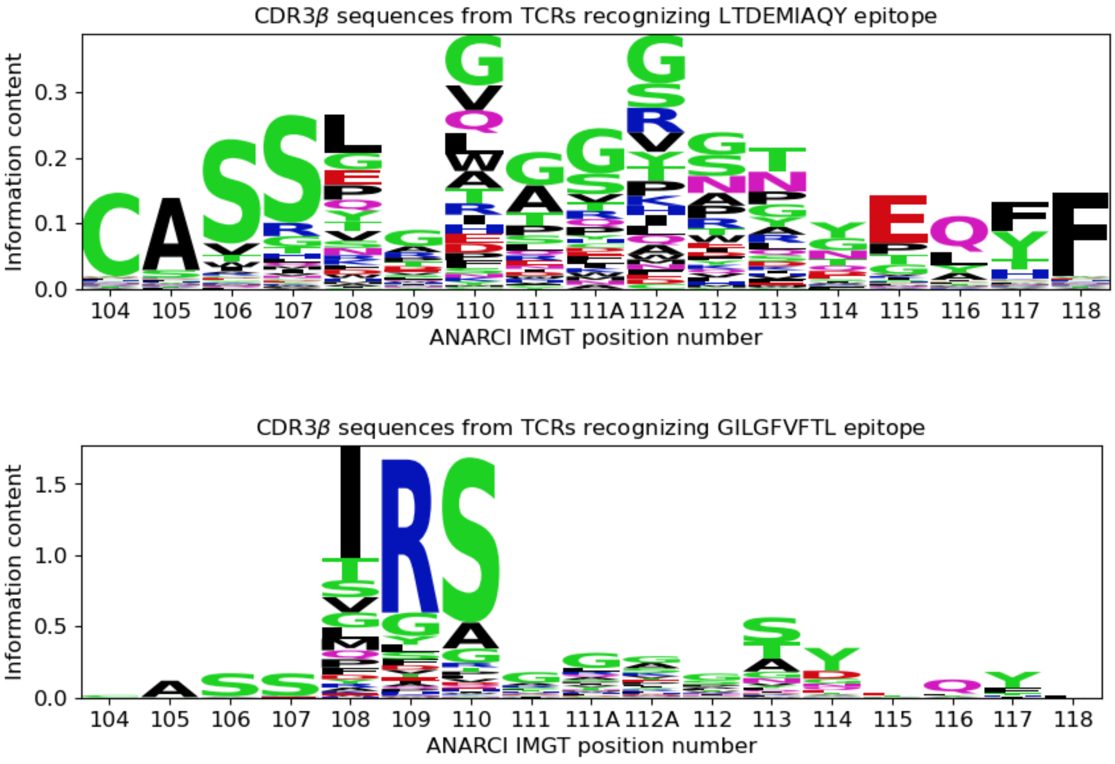


Figure S7. Differential sequence logos of peptide-specific TCRs from IMMREP_2022 data sets - CDR3$\beta$

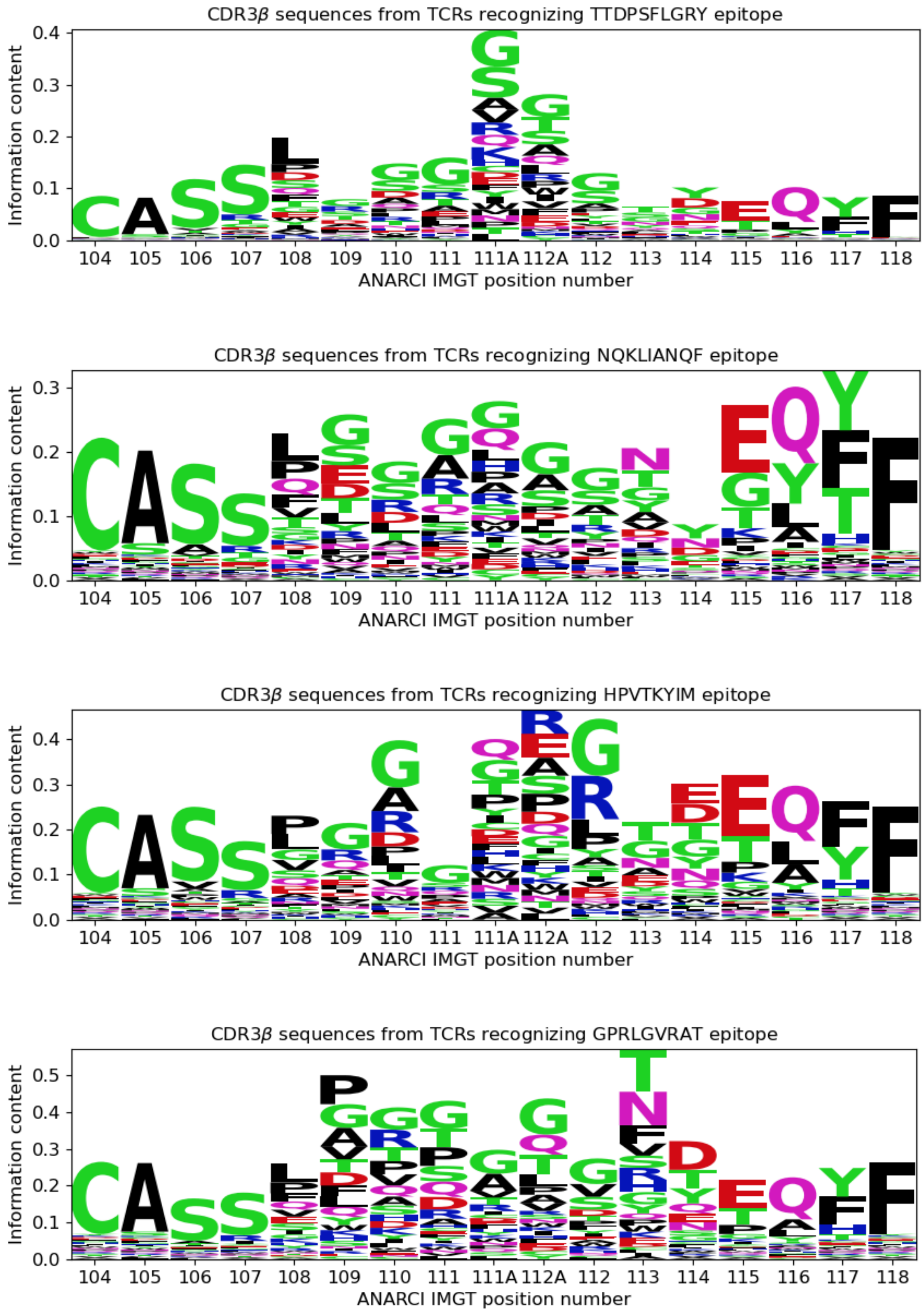


Figure S7. Differential sequence logos of peptide-specific TCRs from IMMREP_2022 data sets - CDR3$\beta$ (continued)

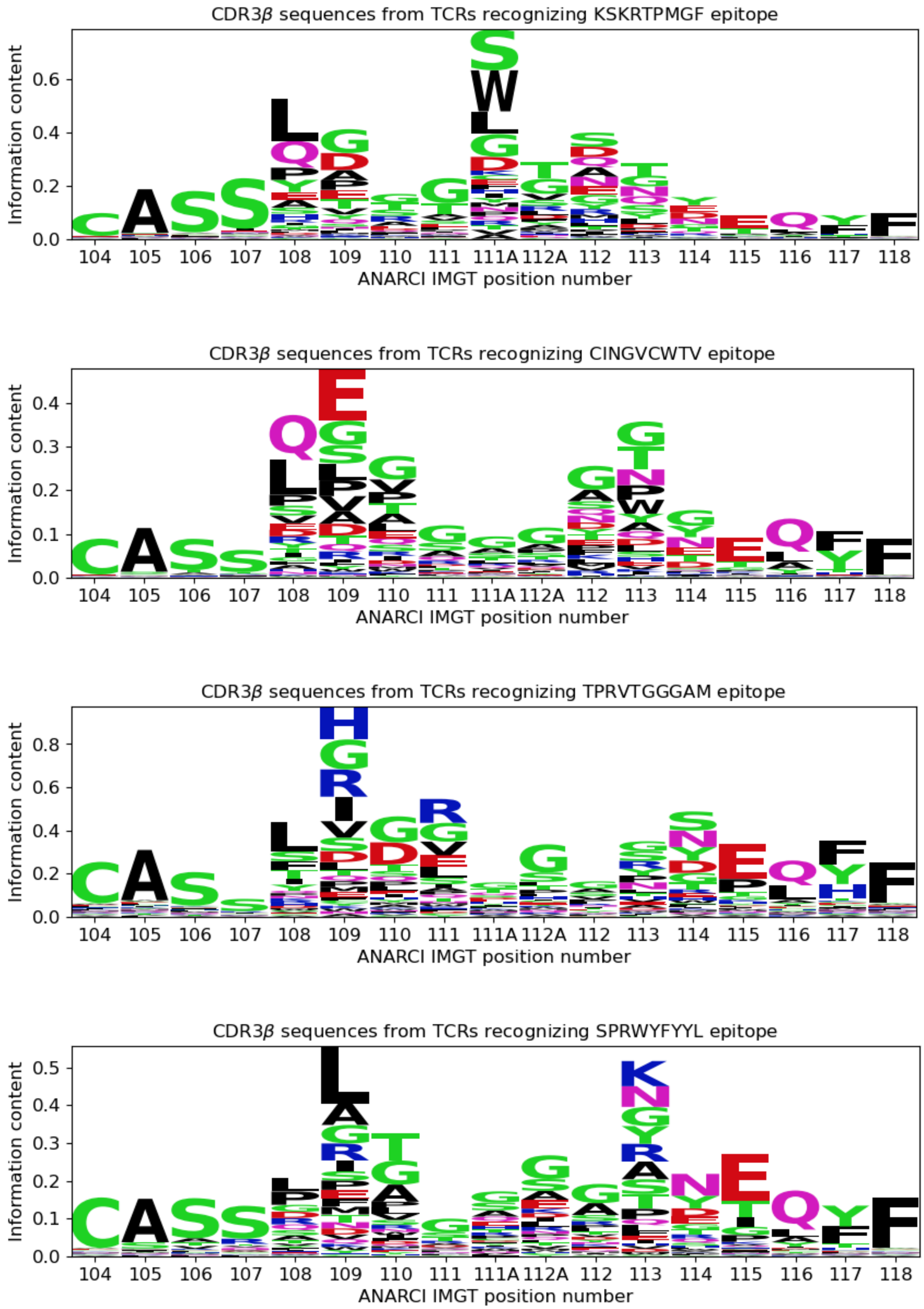


Figure S7. Differential sequence logos of peptide-specific TCRs from IMMREP_2022 data sets - CDR3$\beta$ (continued)

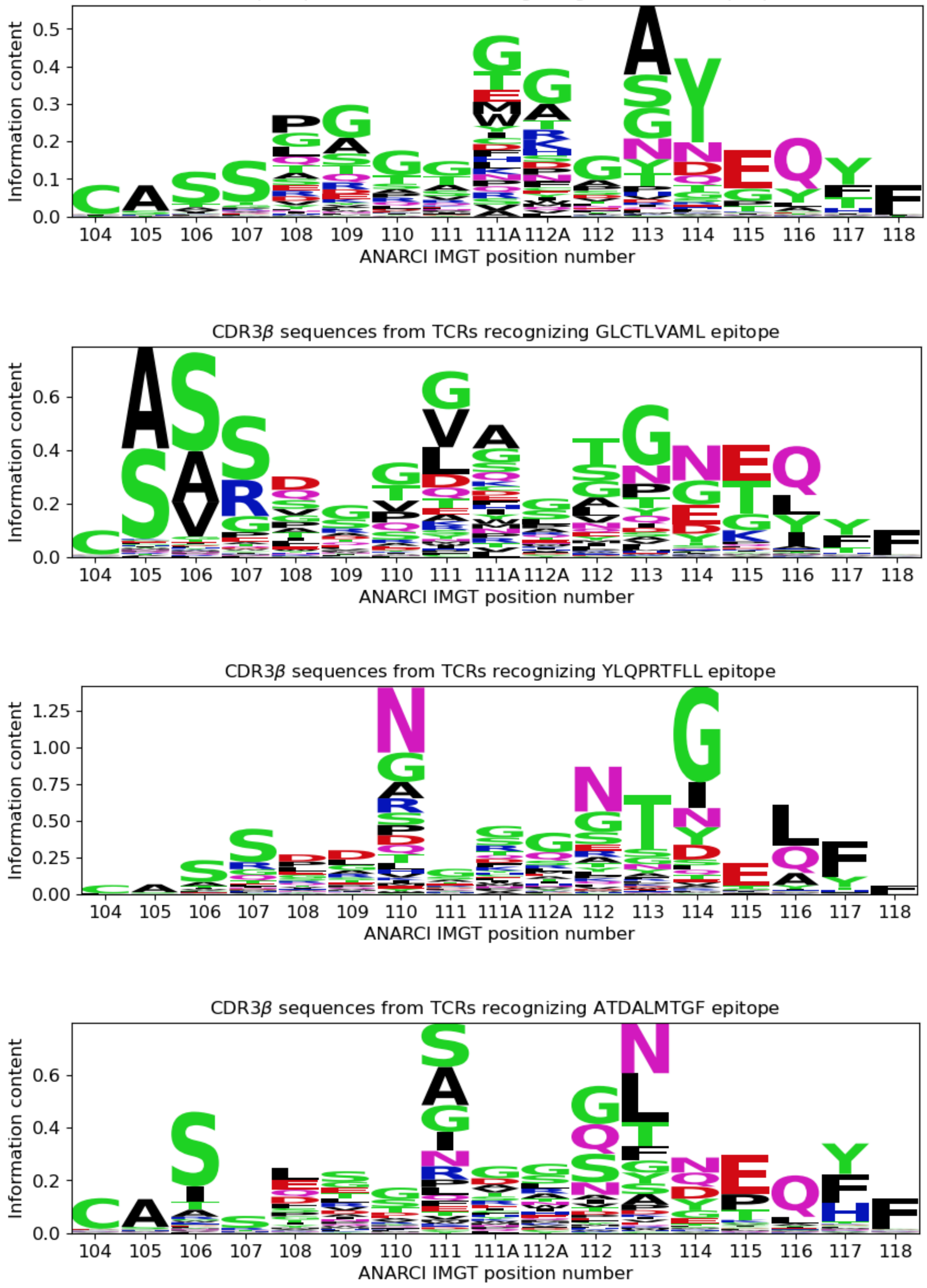


Figure S7. Differential sequence logos of peptide-specific TCRs from IMMREP_2022 data sets - CDR3$\beta$ (continued)

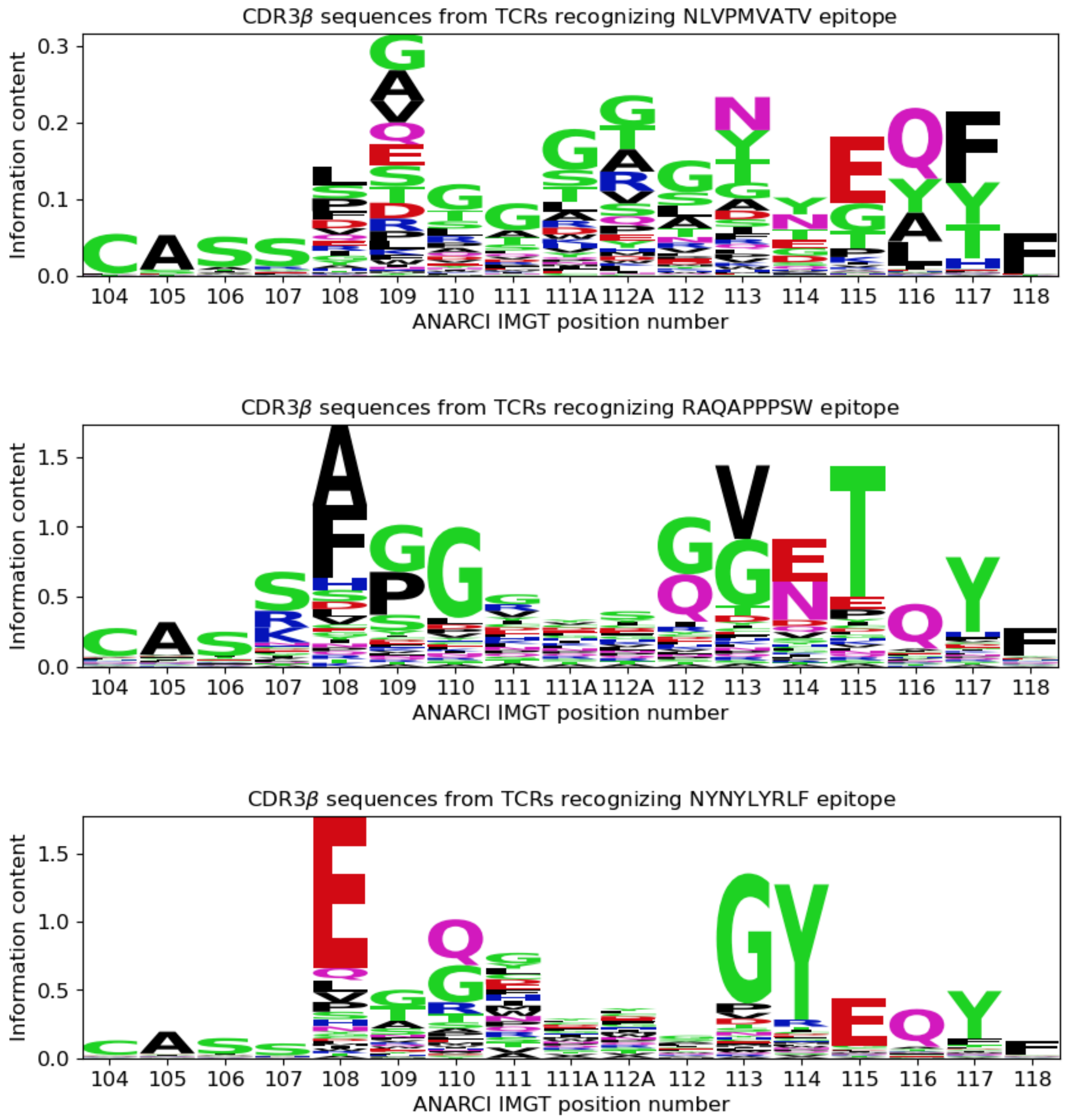


Figure S7. Differential sequence logos of peptide-specific TCRs from IMMREP_2022 data sets - CDR3$\beta$ (continued)

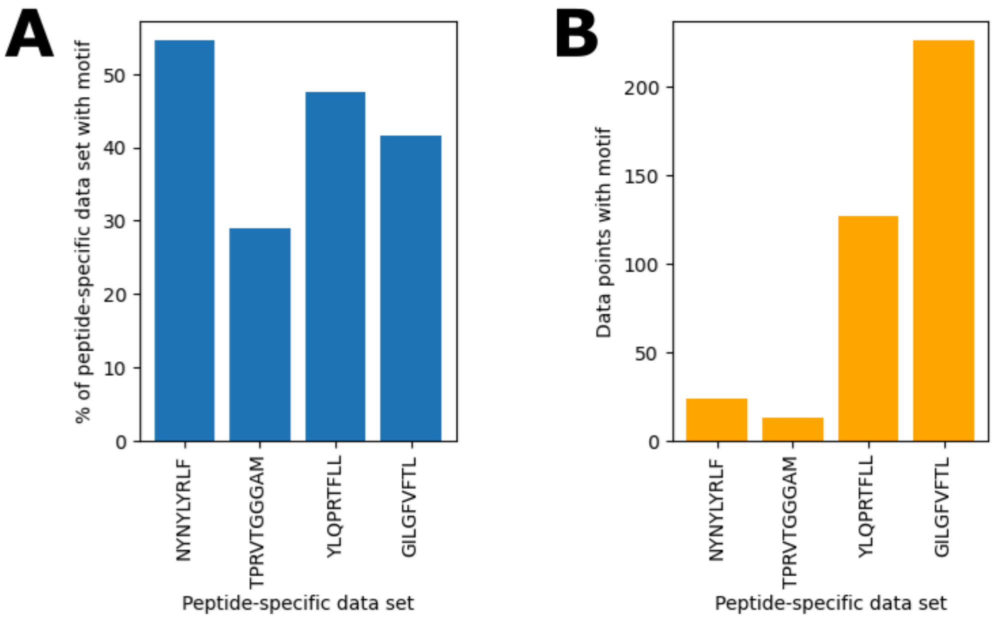


Figure S8. Per-peptide statistics of data points with sequence motifs. (**A**) Percentage in each peptide-specific data set, (**B**) Absolute data point counts.

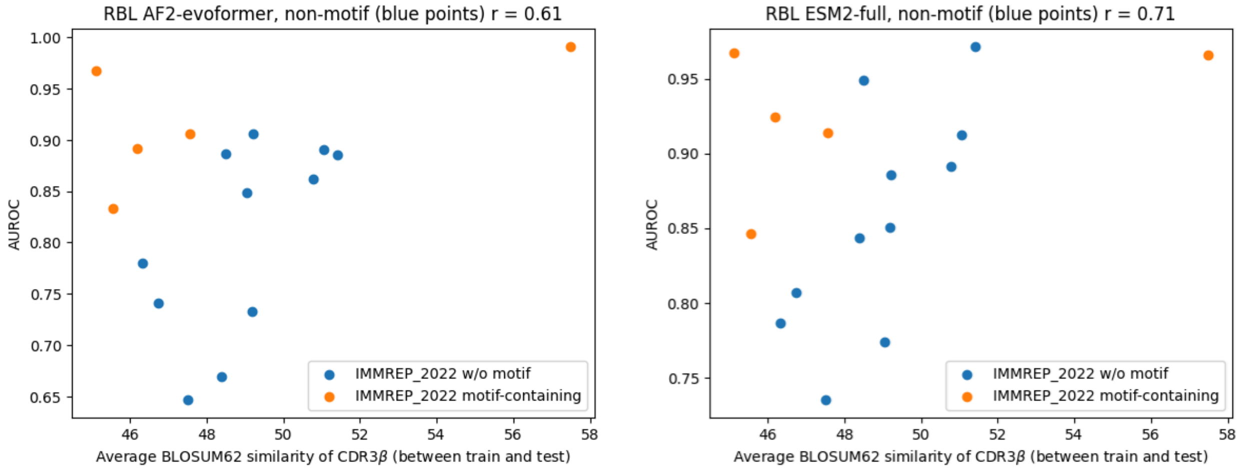


Figure S9. Relation between peptide-specific model performance and average similarity between train and test TCRs (CDR3b sequences). Motif-containing data points are clear outliers.

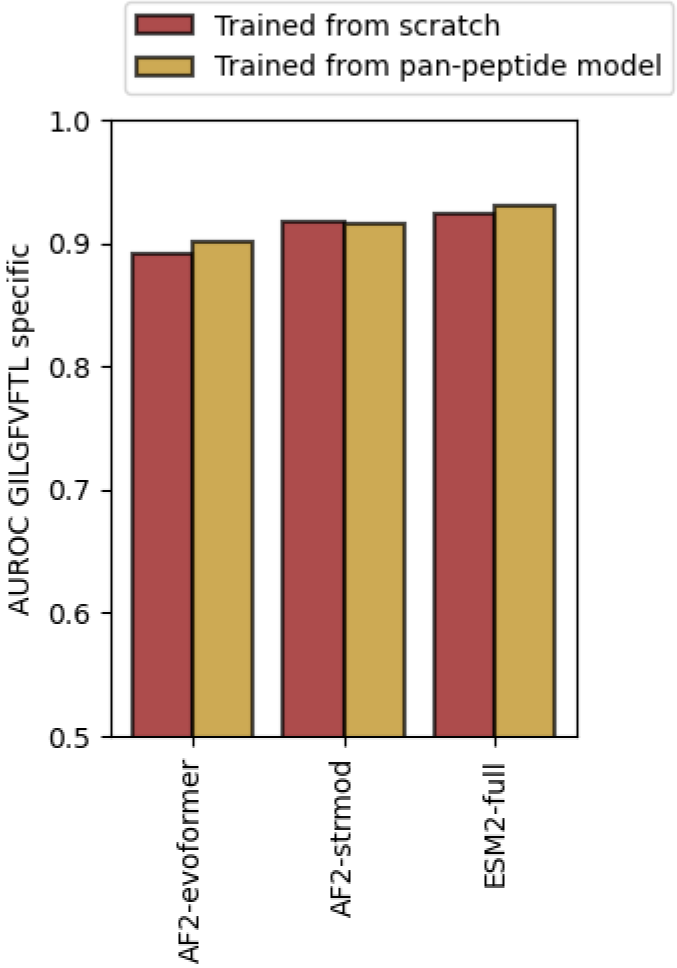


FIgure S10. Effect of using pre-trained pan-peptide models as a starting point for training of various TCRcube models in the GILGFVTL-epitope-specific data set. Note that the pan-peptide model was trained on a modified train set with GILGFVTL excluded.
